# Transcription factor co-occupancy shapes the somatic mutational landscapes across regulatory regions in liver cancer

**DOI:** 10.64898/2026.08.22.745658

**Authors:** Nija George, Anurag Kumar Singh, Radhakrishnan Sabarinathan

**Affiliations:** National Centre for Biological Sciences, Tata Institute of Fundamental Research, Bengaluru, 560065, India

## Abstract

Both global and local chromatin structure influence the heterogeneous distribution of somatic mutations across the cancer genome. For instance, at the local scale, DNA regions bound by transcription factors (TF) and other chromatin-associated proteins display elevated somatic mutation rates, due to variable DNA damage and repair at these protein-bound sites. However, the contribution of TF co-binding towards the variations in somatic mutation rates remains largely unexplored. Here, we combine somatic mutations from whole-genome sequencing of liver cancers with ChIP-seq profiles for over 150 TFs and chromatin-associated proteins in the human liver cancer cell line HepG2, to systematically examine how TF co-occupancy shapes local mutational landscapes. We show that somatic mutation rates at binding sites vary substantially across distinct TFs and co-binding combinations. Furthermore, the magnitude and spatial distribution of somatic mutation rates at TF binding sites differ across promoters and enhancers, likely influenced by the local chromatin accessibility and architecture. Finally, we identify NFIA (Nuclear Factor IA) as a distinct exception, maintaining elevated mutation rates across its binding sites independent of local co-binding context. Together, these findings reveal that combinatorial TF co-occupancy and local chromatin architecture are associated with differences in somatic mutation rates across regulatory regions in liver cancer.

## Introduction

DNA is continually subjected to a wide variety of lesions arising from endogenous cellular processes or exogenous mutagens^1,2^. While cellular repair pathways resolve most DNA damage, unrepaired lesions can give rise to mutations during DNA replication or through error-prone DNA repair^1,3^. Mutations arising in germ cells contribute to heritable genetic variation^4^, whereas mutations occurring in somatic cells accumulate throughout an individual’s lifetime and are not inherited^5^. However, some of these somatic mutations can contribute to clonal expansion in both normal and malignant tissues^5–7^. Across malignancies, somatic mutation burdens vary widely^8–10^, from highly mutated cancers such as melanoma^11^ to low-burden paediatric tumours^12^. Within cancer genomes, the spatial distribution of somatic mutations is also highly heterogeneous. At the megabase scale, this variation is shaped by cell lineage, chromatin compactness, replication timing, and transcriptional activity^9,13–16^. At the local scale, sequence context and chromatin-associated factors, such as nucleosomes and bound transcription factors (TFs), further modulate mutational heterogeneity across the genome^1,17,18^.

For example, transcription factor binding sites (TFBS) exhibit localised increases in somatic mutation density in melanoma and lung cancers, which has been associated with impaired nucleotide excision repair (NER) at protein-bound DNA^17,19,20^. Similarly, lineage-specific TF binding has been associated with altered DNA repair: in prostate cancer, somatic mutations disproportionately accumulate at androgen receptor (AR) bound regions, with AR occupancy interfering with base excision repair (BER) of abasic sites^21^. In breast cancer, estrogen receptor (ER) binding sites showed an elevated somatic mutation rate, capable of altering long-range chromatin contacts between regulatory regions^22^. In hepatocellular carcinoma (HCC), several TFBS have been shown to have an increased mutation rate^23^. In addition to these tumour type-specific patterns, the DNA regions bound by chromatin-architectural proteins such as CTCF and cohesin, at the boundaries of topologically associating domains, exhibit elevated somatic mutations across multiple cancer types^24^. Our recent work showed that this enrichment is largely attributable to the replication-associated DNA damage and impaired repair at CTCF/ cohesin-bound regions^25^. Together, these studies suggest that local mutation patterns at protein-bound DNA vary with distinct TF identities and cellular contexts, with differences in DNA damage formation and repair contributing to this variation^19,26^.

However, the above studies were largely focused on individual TFs or selected TF families, limiting systematic comparisons across many TFs within the same cellular context^17,22^. Moreover, how differences in TF occupancy and multi-factor co-binding relate to the mutational landscape of regulatory regions has not yet been systematically examined. To address this, we leveraged the ChIP-seq profiles available (from ENCODE) in a single liver cancer cell line, HepG2, comprising 208 TFs, cofactors, and chromatin regulators^28,29^. For simplicity, we refer to the analysed binding sites of both TFs and other chromatin-associated proteins collectively as TFBS throughout the manuscript. By integrating whole-genome somatic mutations from hepatocellular carcinoma (HCC) cohort from the Pan-Cancer Analysis of Whole Genomes (PCAWG)^10^ with the ChIP-seq profiles, we examined how TF binding, co-occupancy, local accessibility, and 3D chromatin topology are associated with somatic mutation patterns across regulatory DNA. We focused on liver cancer because it provides a biologically relevant tissue context for this analysis and is characterised by high rates of somatic mutation in both normal and malignant tissues, reflecting the combined effects of diverse endogenous and exposure-associated mutational processes^6,27^. Our analysis revealed that local mutation enrichment varies markedly across TFBS and co-binding contexts and differs between promoter- and enhancer-associated regulatory regions. We further identify NFIA (Nuclear Factor IA) as a distinct case in which elevated mutation rates persist largely independently of local co-binding context.

## Results

### Elevated somatic mutation profiles at TFBS in HCC

To study the variations in somatic mutation patterns across TFBS in liver cancer, we mapped somatic single-base substitutions (SBS) from the PCAWG^10^ HCC cohort (n = 314) to binding sites for 156 TFs (133 DNA-binding factors and 23 chromatin regulators or co-factors) retained after filtering for peak coverage from the HepG2 ENCODE ChIP-Seq datasets^28,29^ (**Fig. 1A**, Supplementary Table 1; See Methods). Baseline hepatic expression of the above TFs was verified using RNA-seq data from the TCGA-LIHC and GTEx liver cohorts^30^ (**Supplementary Fig. S1A**). Somatic mutations were analysed across ±1 kb (2,001-bp) around each ChIP-seq peak midpoint to compare the mutation rate at TFBS with its immediate flanking regions (as local genomic control). Further, to control for the effect of local sequence context, we calculated the mutational enrichment (observed/expected mutations; i.e. fold change [FC]) for each TF, focusing on the core TFBS (101 nt; ±50-bp from the ChIP-seq peak midpoint)^26^ (**Fig. 1A and 1B, Supplementary Fig. S1B**; See Methods).

**Figure 1:**
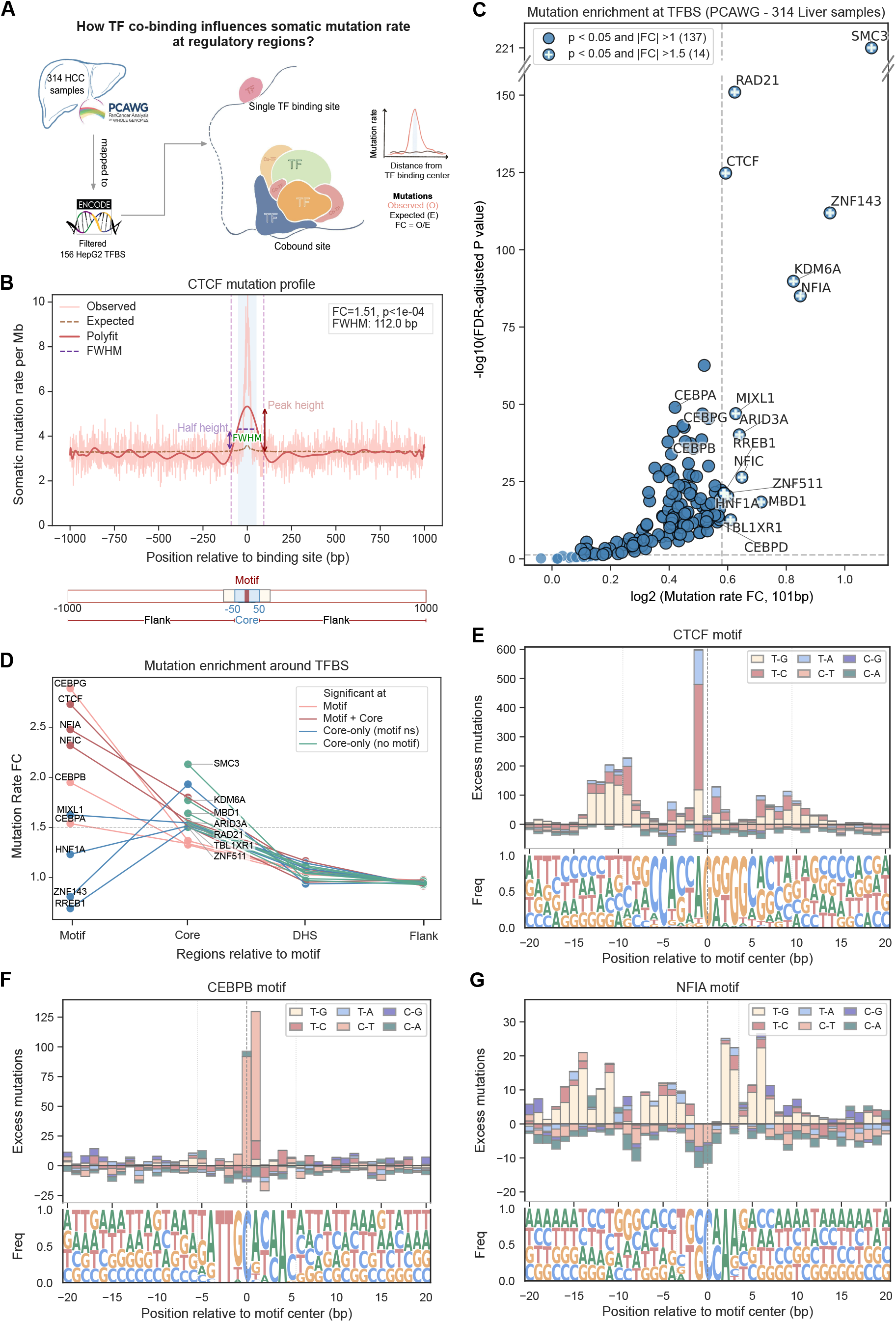
TFBS show elevated somatic mutation profiles in HCC. A. Overview of the study. Single-base substitutions (SBS) from the PCAWG HCC cohort were mapped onto ChIP-seq regions for 156 high-confidence transcription factors (TFs) and chromatin-associated proteins in theHepG2 cell line (obtained from ENCODE). Observed mutation rates were compared with expected rates at individual TF binding sites (TFBS). The mutational rate enrichment fold change (FC) was calculated as the ratio of observed versus expected rates at TF binding sites. Downstream analyses examined the association between mutation rate enrichment and TF co-occupancy. B. Mutation rate profile at CTCF binding sites is shown as a representative example. Somatic mutation rate (per Mb) calculated using the SBS from 314 samples is shown over a 2,001-bp window centred on the CTCF ChIP-seq peak midpoint. FC and statistical significance were calculated using Fisher’s exact test by considering the mutation counts within the central 101-bp core window (±50-bp) with respect to the immediate flanking regions (excluding the core region). The red line shows the smoothed observed mutation rate, and the dashed brown line shows the expected background rate. The horizontal purple dashed line is at the half-maximum level used to calculate the full width at half maximum (FWHM). The schematic below indicates the motif, 101-bp core (−50 to +50-bp) and immediate flanking regions. C. Mutational enrichment across 156 TFBS. Log_2_FC within the 101-bp core region is plotted against −log₁₀(FDR-adjusted P value) for 156 TFBS. The horizontal dashed line marks FDR = 0.05, and the vertical dashed line marks log_2_FC = 0.58 (FC = 1.5). The number of significant TFBS is shown in brackets in the legend. Significant TFBS are outlined in black; significant factors with FC > 1.5 are marked “+”. D. FC across binding-site sub-regions. Comparison of FC across motif, 101-bp core (−50 to +50-bp from motif/ChIP-seq centre), DHS (−250 to 250-bp excluding the core), and distal flanking regions (regions excluding the core and DHS) for TFBS grouped by motif resolution. TFBS showing motif-selective enrichment are shown in orange (CEBP family), core-enriched TFBS with mapped JASPAR motifs in red, and core-enriched TFBS without mapped JASPAR motifs in green. **E-G.** Sequence composition at TF binding sites. The upper panel shows positional excess mutations, calculated as observed minus expected mutations, across pyrimidine-centred substitution classes in a 41-bp window centred on the mapped motif. The lower panel shows the nucleotide frequency logo centred on the TFBS motif midpoint. **E.** Centered on CTCF motif **F.** Centered on CEBPB motif **G.** Centered on NFIA motif.

Consistent with findings in other malignancies^17,24^, we observed a localised accumulation of somatic mutations concentrated at the TFBS core relative to the immediate flanking regions across the majority of the examined TFBS (**Fig. 1B, Supplementary Fig. S1C-R**). In total, 137 of the 156 analysed TFBS (87.8%) showed statistically significant mutational enrichment within the 101-nt core region (two-sided Fisher’s exact test, false discovery rate [FDR] < 0.05) (**Fig. 1C**). Among these, only 14 TFBS displayed strong core mutational enrichment at their respective binding sites with FC exceeding 1.5 (log2FC > 0.58). These 14 significant TFBS represented a functionally highly diverse set of regulators, comprising CTCF and the cohesin subunits RAD21 and SMC3, which govern 3D genome architecture^24,31,32^; essential hepatic regulators like HNF1A, which modulates liver-specific transcriptional programs^33^ and has been implicated in hepatitis B virus (HBV) transcription and replication^34,35^; NFI-family regulators NFIA and NFIC, which participate in developmental, transcriptional and chromatin-regulatory programs^36,37^; early developmental factor MIXL1^38^, transcriptional regulator RREB1^39^ and co-regulator TBL1XR1^40^; and zinc-finger factors ZNF143 and ZNF511, with loss of ZNF143 reported to alter distal regulatory interactions and cohesin occupancy at CTCF-bound sites^41^.

Similar to the PCAWG HCC cohort (**Fig. 1C**), the enrichment of somatic mutations across TFBS was observed in an independent liver cancer cohort (ICGC - LICA-CN^42^), where 130 out of 156 TFBS profiles (83.3%) were significantly enriched in mutations at the core region (**Supplementary Fig. S2A**). Moreover, a non-cancer cohort^6^ exhibited a similar directional pattern concordant with the FCs at TFBS cores for the cancer cohorts, but not statistical significance (at FDR < 0.05) (**Supplementary Fig. S2B**). The stronger statistical signal in HCC may reflect the higher somatic mutation burden and mutational processes associated with malignant liver^6,27^.

### Motif-level resolution reveals distinct mutational patterns

To examine the mutation patterns at higher resolution, we mapped position weight matrices from JASPAR 2024^43^ to the HepG2 ChIP-seq regions using the MOODS suite^44^ and precisely located the binding motif for 90 of the 156 TFBS (**Supplementary Fig. S3A**, Supplementary Table 2; See Methods). Analysis of these motif-centred regions provided nucleotide-level resolution and identified six TFBS (CTCF, NFIA, NFIC, CEBPA, CEBPB and CEBPG) with significantly high mutational enrichment at the motif level (FC > 1.5; **Fig. 1D, Supplementary Fig. S3B**). CTCF, NFIA and NFIC binding sites displayed sharply localised mutational enrichment centred on the motif and 101-nt core, which progressively declined towards the surrounding DNAse Hypersensitivity Sites (DHS) and distal flanking regions (**Fig. 1D, Supplementary Fig. S3D, S1F and S3N**; See Methods). However, this enrichment was absent at TF motifs lacking ChIP-seq signal, indicating that the underlying motif sequence alone is insufficient to explain the elevated mutation rate at bound sites (**Supplementary Fig. S3C**).

The CEBP family (CEBPA, CEBPB and CEBPG), which did not meet the FC > 1.5 threshold at the broader TFBS core level, displayed strong mutational enrichment when centred on their cognate motifs (**Supplementary Fig. S3B, S3E, S3G and S3H**). Conversely, motif regions of core-enriched TFBS HNF1A, MIXL1, ZNF143 and RREB1 exhibited lower mutational burden at the immediate motif relative to the surrounding 101-nt core, followed by a decline as seen for other TFBS, toward the flanking regions **(Fig. 1D, Supplementary Fig. S3I, S3M and S3O**). A similar trend was observed for TFBS lacking mapped motifs (**Fig. 1D**), indicating that the elevated mutational frequency at these regulatory regions is associated with protein occupancy on the DNA, either directly or indirectly.

The elevated mutation burden at TFBS motifs was investigated to check whether specific substitution types were preferentially enriched. Characterisation of the nucleotide substitution classes at the motif-centred regions revealed a prominent excess of observed T>C and T>G substitutions over the expected (**Fig. 1E and 1G, Supplementary Fig. S3**; See Methods). TF binding regions for CTCF revealed highly concentrated T>C/G enrichment at the motif (**Fig. 1E**), with mutation rates differing across positions within the CTCF motif^26,27,45^, whereas the hepatic lineage-specific HNF1A showed a more diffuse distribution of these substitutions across the motif and surrounding TF binding region (**Supplementary Fig. S3L**). In contrast, the CEBPB motif exhibited a distinct, disproportionate accumulation of C>T transitions in a CpG context (**Fig. 1F**), as did CEBPA and CEBPG (**Supplementary Fig. S3J and S3K**), whereas NFIA-associated mutations were characterised by an excess of T>G substitutions and relative depletion of C>T and C>A events across the motif-centred window (**Fig. 1G**). CEBP family members (CEBPA, CEBPB and CEBPG) have established roles in maintaining hepatic transcriptional programs and liver regeneration^46^. The observed enrichment of C>T at the CpG context could be explained by a distinct mutational process (such as deamination of methylated CpGs^47^) along with TF occupancy. Taken together, these results show that local sequences can act jointly with TF occupancy to influence the mutation density in regulatory regions.

### Peak topology distinguishes spatial patterns of mutational enrichment at TFBS

We next asked whether the mutation enrichment, tightly confined to the motif region or extending more broadly into the surrounding (core) region, could be explained by multiple TF co-binding or proximity to the transcription start site (TSS) and associated transcriptional machinery^26^. For this, we quantified the Full Width at Half Maximum (FWHM) for each TFBS mutation profile and compared it with core mutation rate FC (**Fig. 1B**; See Methods). FWHM varied substantially across TFBS profiles (median, 171 nt; interquartile range, 131-199 nt; maximum, 315 nt); however, the TFBS with significant mutation enrichment at the core (FC>1.5) were confined to the range of 100 nt to 200 nt.

The variability in FWHM raises the possibility that proximity to the TSS may contribute to differences in mutation-peak width. We therefore calculated the proportion of mutated TFBS that are further away (distal) from the TSS proximal regions (that is, -2kb to +500 bp from GENCODE-defined TSS^48^; **Fig. 2A**; See Methods). Across the 156 mutation profiles, mutated TFBS were predominantly distal (median distal fraction, 73.8%; interquartile range, 60.4 - 85.5%). TFBS with lower distal fractions (<74%) tended to show lower FWHM (<171 nt) and lower core mutation enrichment (FC < 1.5), whereas many profiles with higher distal fractions (≥ 74%) showed broader peaks (FWHM ≥ 171 nt) but with lower mutation enrichment (FC < 1.5). The lower mutation enrichment at more TSS-proximal TFBS may partly reflect more efficient DNA repair within accessible and transcriptionally active regulatory regions, as reported previously^18,49^. Nevertheless, several TFBS with significant mutation enrichment (FC > 1.5) deviated from this trend. CTCF, SMC3, RAD21 and NFIA were predominantly distal and showed narrow mutation peaks (**Fig. 2A, Supplementary Fig. S1K, S1M, S1O**>), whereas NFIC was similarly distal but showed a broader profile (**Fig. 2A, Supplementary Fig. S1L**). Thus, TSS proximity alone did not explain the differences in mutation-peak width and enrichment across TFBS, prompting us to examine if local TFBS co-binding contributed to these distinct profiles.

**Figure 2:**
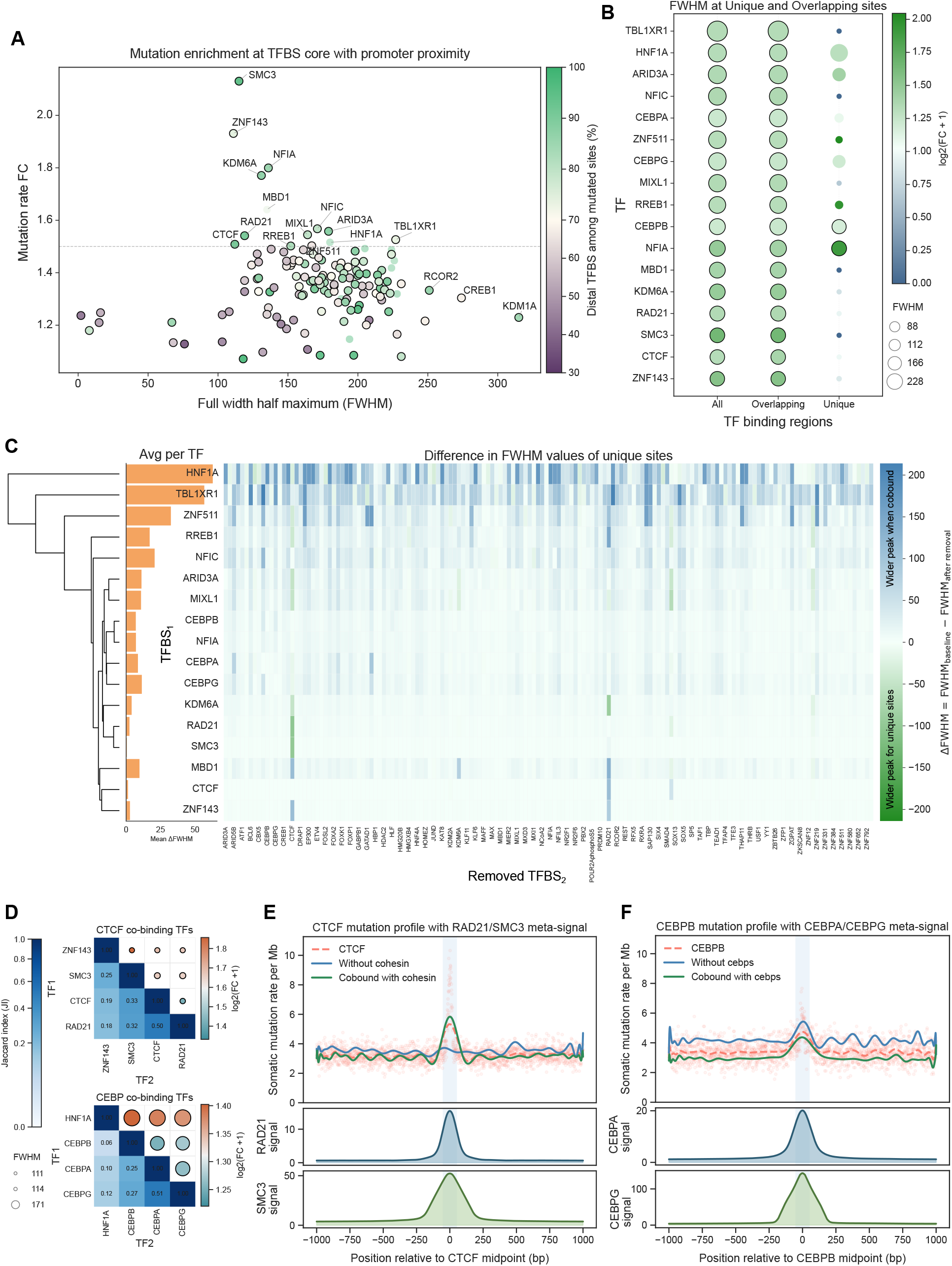
TF co-binding is associated with mutational enrichment and footprint width at TFBS. **A.** Relationship between mutation enrichment and peak width. Core FC is plotted against FWHM to distinguish relatively narrow and broad mutation profiles. Point colour indicates the fraction of mutated TFBS located distal to the nearest transcription start site (TSS), with proximal sites defined as −2,000 to +500-bp relative to the TSS. Enrichment or depletion of mutations at proximal versus distal TFBS was assessed using Fisher’s exact test by comparing mutated and non-mutated TFBS. Points with significant (FDR < 0.05) proximal/ distal associations are outlined in black. **B.** FC and FWHM across co-binding categories. Bubble plot showing mutational FC in the 101-bp core window for 17 prioritised TFBS profiles. TFBS are partitioned into all sites, sites overlapping at least one of the other 155 TFBS sets, and non-overlapping (unique) sites without overlap with any of the other TFBS sets. Bubble size indicates peak span, measured as FWHM. TFBS with significant mutation enrichment (FDR < 0.05) are outlined in black. **C.** ΔFWHM after removing co-bound partner TFs. Hierarchically clustered heatmap showing changes in FWHM after sequentially removing sites overlapping each of the other 155 TFBS sets from each of the 17 prioritised profiles. ΔFWHM is defined as FWHM_baseline_ − FWHM_after removal_. Positive values (blue) indicate peak narrowing after removal, values near zero indicate little change, and negative values (green) indicate peak broadening after removal. The bar plot on the left shows the mean ΔFWHM for each of the prioritised TFBS. **D.** Co-occupancy and mutational dependency between TFs. Composite matrices showing TF co-occupancy, measured by Jaccard index (JI) in the lower triangle, and FC and FWHM after partner-specific overlap partitioning in the upper triangle. Results are shown for the CTCF–cohesin module in the upper panel, including CTCF, SMC3 and RAD21, with ZNF143 shown for comparison, and the CEBP family module in the lower panel, including CEBPB, CEBPA and CEBPG, with HNF1A shown for comparison. **E.** Mutational footprints and cohesin occupancy at CTCF binding sites. The upper panel shows somatic mutation rate (per Mb) across a 2,001-bp window at all CTCF binding sites (FC = 1.51, p < 1e-04), CTCF sites remaining after removal of sites overlapping RAD21 or SMC3 (FC = 0.98, p = 0.56), and CTCF TFBS overlapping RAD21 or SMC3 (FC = 1.68, p < 1e-04). The lower panel shows RAD21 and SMC3 ChIP-seq coverage across the same CTCF-centred window. **F.** Mutational footprints and family member occupancy at CEBPB binding sites. The upper panel shows mutation rate profiles across a 2,001-bp window at all CEBPB binding sites (FC = 1.33, p < 1e-04), CEBPB sites remaining after removal of sites overlapping CEBPA or CEBPG (FC = 1.25, p < 1e-04), and CEBPB TFBS overlapping CEBPA or CEBPG (FC = 1.38, p < 1e-04). The lower panel shows CEBPA and CEBPG ChIP-seq coverage across the same CEBPB-centred window.

### TF co-occupancy is associated with somatic mutation accumulation at regulatory DNA

Given that regulatory proteins frequently occupy shared genomic regions where multiple factors can influence local DNA accessibility and chromatin conformation^28,50^, we examined whether the localised mutation enrichment observed at the 17 prioritised TFBS was associated with TF co-binding. These comprised the 14 TFBS with core FC > 1.5 together with the three additional motif-enriched CEBP family members (CEBPA, CEBPB and CEBPG) (**Fig 1D**). Co-occupancy was defined by overlap between ChIP-seq-defined binding intervals in HepG2, which could indicate but not necessitate simultaneous binding within individual cells^28^. For each of the 17 TFBS profiles, we partitioned binding intervals into overlapping sites (co-bound by or intersecting with at least one of the remaining 155 TFBS sets) and non-overlapping or unique sites (bound by the target TF alone and no overlap with any of the other; See Methods). At overlapping sites, observed mutation rate profiles, core FC, and peak widths (FWHM) closely resembled the patterns observed at the original unpartitioned TFBS sets (**Fig. 2B**). At unique TFBS, a reversion to expected mutational enrichment was observed, accompanied by substantial changes in peak shape for most profiles (**Fig. 2B, Supplementary Fig. S4C**). Thus, much of the strong enrichment observed in the original TFBS profiles was concentrated at genomic regions shared with other ChIP-seq-defined binding sites. NFIA and CEBPB were notable exceptions. Both retained significant core mutation enrichment at their unique, non-overlapping genomic intervals, with FCs of 2.74 (baseline unpartitioned TFBS FC = 1.80) and 1.26 (baseline unpartitioned TFBS FC = 1.37), respectively (**Fig. 2B**). This indicates that while co-occupancy is associated with mutation enrichment across most regulatory regions, NFIA and CEBPB retain mutation abundance independent of their co-binding context.

To determine if the observed alterations were driven broadly by many overlapping TFBS or selectively by particular partners, we next performed an iterative leave-one-out subtraction analysis. For each of the 17 TFBS, we sequentially removed sites overlapping each of the remaining 155 TFBS and recalculated the parameters of the mutation profiles. The resulting shift in peak geometry was defined as ΔFWHM (FWHM_baseline_ - FWHM_after removal_; Fig. 2C). A positive ΔFWHM indicates narrowing of the mutation peak after removal of sites overlapping a given partner, values near zero indicate little change, and negative values indicate broadening after removal. Hierarchical clustering of these ΔFWHM profiles revealed three broad co-binding dependency classes: broad sensitivity, selective sensitivity and resilience to removal of overlapping TFBS (**Fig. 2C**). Within the broadly sensitive groups, HNF1A, TBL1XR1, ZNF511, RREB1 and NFIC showed pronounced changes in peak width and loss of core enrichment following removal of sites overlapping many different TFBS sets whereas, ARID3A, MIXL1, MBD1, CEBPA, and CEBPG portrayed moderate changes following removal of overlapping sites of individual TFBS sets. This broad sensitivity may indicate that these TFBS may occur within dense multi-factor hubs or regulatory regions whose mutational patterns depend on the surrounding TF density. Among the selectively sensitive TFBS, CTCF and the cohesin-associated factors RAD21 and SMC3 displayed a distinct interdependent pattern. Mutation rates and peak geometries at CTCF sites collapsed specifically when sites overlapping RAD21 or SMC3 were excluded, but remained largely resilient to the removal of unrelated TFs. Smaller changes were also observed after removing CTCF sites overlapping KDM6A, CEBPG and ZNF143. This selective pattern suggests that the CTCF mutational footprint is particularly concentrated at sites shared with cohesin rather than being equally dependent on overlap with all TFBS. The resilient group comprised NFIA and CEBPB. Confirming our unique-site observations, NFIA and CEBPB displayed minimal shifts in ΔFWHM regardless of which individual overlapping TFBS set was removed (**Fig. 2C**). The above observation is consistent with their mutational landscapes being less dependent on detectable local co-binding than those of the other prioritised TFBS.

### TFBS overlap reveals distinct co-binding patterns

The distinct leave-one-out responses suggested that differences in TFBS mutation profiles may be related to the extent of overlap with other TFBS. To study this independently, we calculated pairwise binding site overlap matrices across all 156 TFs using the Jaccard Index (JI; See Methods). Hierarchical clustering of the resulting similarity matrix revealed well-defined spatial co-occupancy modules in HepG2 cells (**Fig. 2D; Supplementary Fig. S4A**).

As expected, CTCF, RAD21, and SMC3 formed a prominent and correlated co-binding module (JI of CTCF with RAD21 = 0.50 and SMC3 = 0.33), reflecting their shared occupancy at architectural loop anchors^24,28,31^ (**Fig. 2D**). ChIP-seq signal coverage profiles confirmed co-localisation of RAD21 and SMC3 centred precisely around CTCF binding midpoints (**Fig. 2E**). Excluding any one of the above cohesin components, resulted in a reduction in core mutation FC (**Fig. 2C and 2E**). This agrees with previous reports of elevated mutation accumulation at CTCF/cohesin co-bound sites^24,51^, and our observations demonstrate that the strongest CTCF-associated mutational footprint occurs at genomic sites with detectable CTCF/cohesin overlap. A second distinct co-binding module was formed by the CEBP family (CEBPA, CEBPB, and CEBPG; **Fig. 2D, Supplementary Fig. S4A**). Unlike CTCF, excluding CEBPB sites overlapping other CEBP family members did not ablate the core mutational footprint of CEBPB (**Fig. 2F**). Previous work has also revealed that C/EBP proteins can preferentially bind G:T mismatches and interfere with their repair, providing a possible autonomous, TF-specific mechanism for mutation accumulation at C/EBP binding sites^47^. The other persistent, mutation-accumulating TF, NFIA, exhibited negligible spatial overlap with all other examined TFs ( **Supplementary Fig. S4A**) while retaining strong core mutational enrichment at unique NFIA binding sites (FC > 2.0; **Supplementary Fig. S4B**). Thus, the spatial organisation of TFBS can underlie mutational dependence on co-binding, but this relationship is not universal, and mutation enrichment can persist irrespective of local TFBS overlap.

### Local epigenomic architecture shows active NFIA binding regions

Given this distinct pattern at NFIA binding sites, we checked whether their persistent mutational enrichment (FC = 1.8; **Fig. 3A**) occurred within an active regulatory chromatin context or could instead reflect accumulation in relatively inactive regions. Using ChIP-seq profiles of various activating and repressing histone marks from the ENCODE portal^29^ (See Methods), we found regions around the NFIA binding midpoint significantly enriched for the active histone modifications H3K27ac, H3K9ac, H3K4me1 and H3K4me3 relative to the local surrounding chromatin (**Fig. 3B**, Supplementary Table 3; See Methods). In contrast, the repressive marks H3K27me3 and H3K9me3 showed substantially weaker local changes. The enrichment of H3K4me1 together with H3K27ac is consistent with enhancer-associated active chromatin, while concurrent H3K4me3 enrichment suggests that NFIA sites span broader active regulatory contexts^52,53^. We also observed prominent bidirectional nascent transcription profiles (GRO-seq^54^) at both mutated and non-mutated NFIA binding sites, indicating that these loci represent active regulatory regions^55^ (**Fig. 3C**; See methods). Consistent with this local chromatin architecture, HepG2 MNase-seq^56^ exhibited reduced nucleosome dyad density at the NFIA binding midpoint, with periodic nucleosome positioning in the flanking regions (**Fig. 3E**). Together, these features support an accessible and transcriptionally active regulatory environment around NFIA binding sites in HepG2. At the same HepG2-defined genomic coordinates of NFIA binding sites, K562 (human leukaemia cell line) and GM12878 (normal B-lymphoblastoid cell line)^29^ displayed greater central nucleosome occupancy accessibility, indicating cell-type-specific differences in binding and nucleosome organisation around these loci (**Fig. 3E**). This suggests that the open chromatin configuration observed at NFIA sites in HepG2 may not be maintained in other cellular contexts.

**Figure 3:**
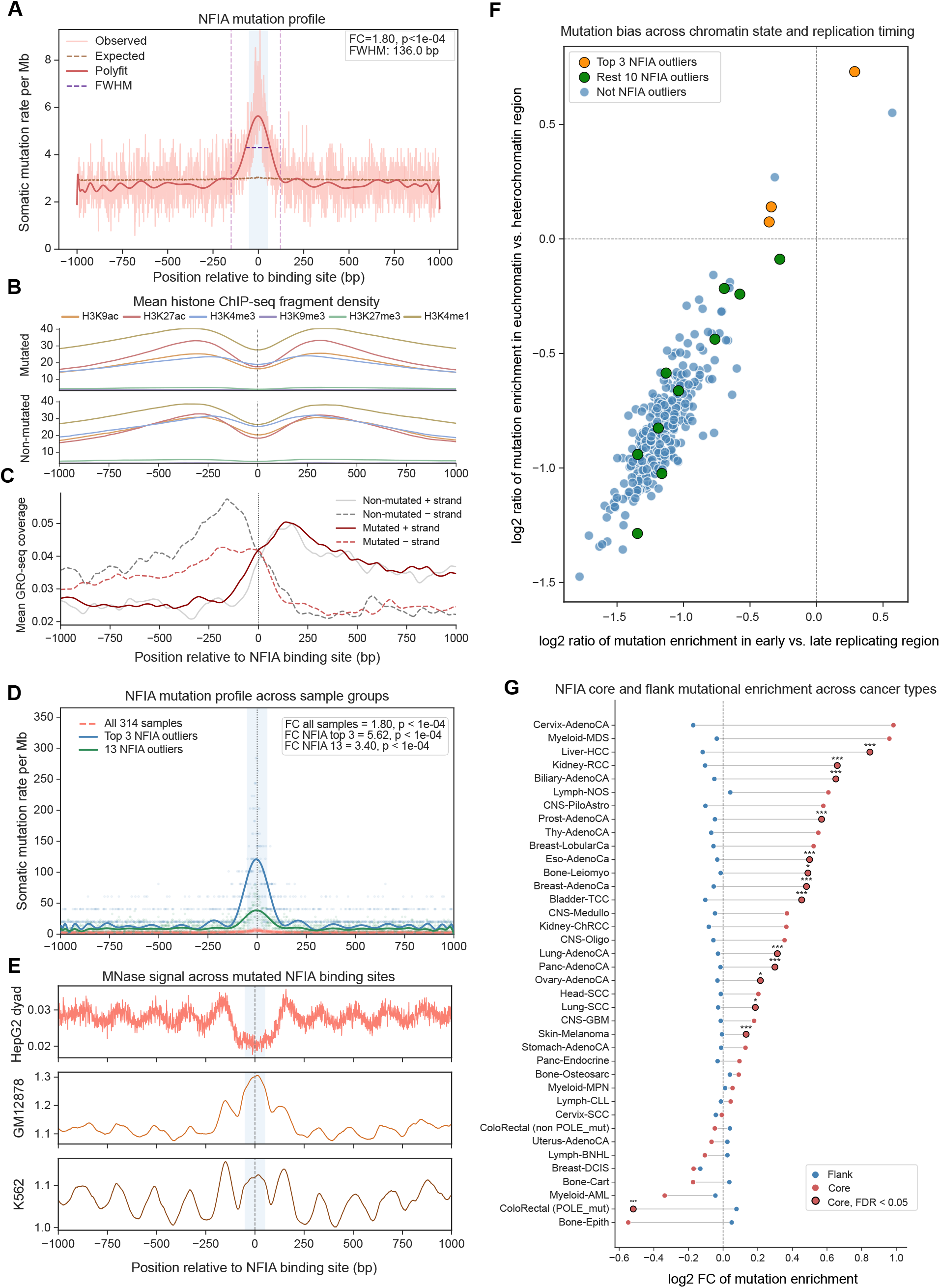
Chromatin context, transcriptional activity, and mutational drivers at NFIA binding sites. **A.** Mutation profile at NFIA binding sites across a 2,001-bp window. Somatic mutation rate (per Mb) calculated from 314 samples is shown over a 2,001-bp window centred on the ChIP-seq peak midpoint. FC and statistical significance (Fisher’s exact test) were calculated within the central 101-bp core window (±50-bp) with respect to the immediate flanking regions (excluding the core region). The red line shows the smoothed observed mutation rate and the dashed brown line shows the expected background rate. The horizontal purple dashed line marks the FWHM. **B.** Histone mark distribution around NFIA binding sites. Mean ChIP-seq signal intensity for activating and repressive histone modifications is shown across the 2,001-bp NFIA-centred window in HepG2 cells. **C.** Transcriptional activity at mutated and non-mutated NFIA binding sites. Strand-separated GRO-seq signal intensity is shown across the 2,001-bp NFIA-centred window in HepG2 cells, comparing mutated and non-mutated sites. **D.** NFIA mutation profiles across sample groups. Smoothed somatic mutation profiles are shown across a 2,001-bp window centred on NFIA binding sites for all 314 PCAWG samples (orange), the 13 NFIA cohort outliers (green), and the top three NFIA cohort outliers (blue). FC and statistical significance (Fisher’s exact test) were calculated within the central 101-bp core window (±50-bp) with respect to the immediate flanking regions (excluding the core region). **E.** Nucleosome organisation around NFIA binding sites. HepG2 MNase-seq dyad density (top), and MNase-seq nucleosome-density signals from K562 (middle) and GM12878 (bottom) are shown across 2,001-bp windows centred on HepG2-defined NFIA binding sites. **F.** Sample-level mutation distribution by chromatin state and replication timing. Sample-wise log_2_ ratio of observed versus expected mutation enrichment in euchromatin (active) versus heterochromatin (repressed) regions (y-axis), and in early versus late replicating regions (x-axis) in HepG2 cells for the PCAWG cohort. Dashed lines mark the baseline level with no relative bias (log_2_ratio = 0). Highlighted points indicate sample outliers. **G.** Pan-cancer mutational enrichment at HepG2-defined NFIA binding sites. Somatic substitutions from individual PCAWG cancer cohorts were mapped to NFIA binding intervals defined in HepG2 cells. Log_2_FC is shown for the central 101-bp core and flanking regions for each cancer cohort. Significant enrichment or depletion at core determined by Fisher’s exact test is denoted as *P < 0.05, **P < 0.01 and ***P < 0.001.

### Sample-level analysis identifies NFIA mutation-burden outliers

We next asked whether the mutational burden at NFIA binding sites was skewed by a subset of hypermutated tumour samples. Interquartile range (IQR) outlier filtering (Q3 + 1.5 x IQR) across the cohort identified eight cohort-wide hypermutated samples based on total mutation burden and 13 NFIA-specific outlier samples that contributed disproportionately to the mutation burden at NFIA binding sites. Five of the 13 NFIA outliers overlapped the cohort-wide hypermutated group (Supplementary Table 4; See Methods). These 13 samples also exhibited elevated mutation counts across all 17 prioritised TFBS (**Supplementary Fig. S5A**), suggesting that their elevated burden at NFIA binding sites may reflect a broader sample-level mutational phenotype.

By reconstructing the NFIA mutation profile using only the top three and top 13 outlier samples, we tested the sensitivity of our findings to these sample-level biases. While these outliers generated elevated fold changes (FC = 5.62 for top 3; FC = 3.40 for top 13; **Fig. 3D**), confirming that these samples substantially amplify the NFIA signal, removing all 13 outliers from the full cohort did not abolish the mutational footprint. The remaining NFIA non-outlier samples cohort (n = 301) retained significant core mutational enrichment at the NFIA binding site (FC = 1.51), and the global enrichment landscape across all 156 TFs was largely preserved (**Supplementary Fig. S5B**). This suggests that these outlier samples can amplify local mutation counts but are not sufficient to explain the NFIA-associated enrichment observed across the HCC cohort.

### NFIA outlier samples show altered genome-wide chromatin and replication-timing biases

The 13 NFIA outlier samples were further examined to determine if their elevated mutation burden stems from local or genome-wide mutational processes by mapping the somatic mutations across global chromatin states and replication timing domains. Using ChromHMM 15-state annotations from the NIH Roadmap Epigenomics Project^53,57^ and HepG2 Repli-seq datasets^29,58^, we calculated active-to-repressed (euchromatic-to-heterochromatic) chromatin and early-to-late replication observed by expected (O/E) ratios for each sample (**Fig. 3F, Supplementary Fig. S5C and S5D;** See Methods). Early-replicating and euchromatic regions generally accumulate fewer mutations than late-replicating and heterochromatic regions, reflecting regional differences in chromatin organisation, replication timing and DNA repair^13,15,16^. Consistent with this pattern, most HCC samples displayed relatively greater mutation accumulation in late-replicating and heterochromatic genomic regions. In contrast, the top three NFIA outlier samples exhibited positive log_2_(euchromatin/heterochromatin) enrichment ratios and preferential mutation accumulation within early-replicating genomic domains (**Fig. 3F, Supplementary Fig. S5C and S5D**). Therefore, the shift towards euchromatic and early-replicating regions indicates that a subset of NFIA outlier samples reveals global changes in the distribution of mutation across the genome.

A systematic screen of clinical, viral and genomic variables was performed to determine if the broader mutational shift in NFIA outlier samples could be attributed to a shared clinical or mutational process. Donor age (**Supplementary Fig. S6A**) and the effect of hepatitis virus (HBV/HCV) infection status on NFIA expression among samples with both viral and expression data available (**Supplementary Fig. S6B**; See Methods) did not distinguish the NFIA outlier cohort. Age-associated signatures SBS5 and SBS40^2^ contributed largely across the cohort, in both outliers and non-outliers (**Supplementary Fig. S6D**), whereas mismatch repair-associated signatures SBS6 and SBS26^2^ were detected in only two NFIA outliers (Supplementary Table 5). Driver gene analysis further revealed recurrent CTNNB1 locus gains/amplifications in 7 of the 13 evaluated outlier samples (**Supplementary Fig. S6E and S6F;** See Methods). CTNNB1 copy-number gains have been reported to increase β-catenin signalling in HCC^59^, although whether this contributes to the NFIA outlier phenotype remains unclear. The outlier group also displayed a trend towards reduced overall survival (**Supplementary Fig. S6C**), but the small number of samples did not allow us to directly relate this to CTNNB1 copy-number status. These results reveal a shift towards euchromatic and early-replicating regions in a subset of NFIA outliers, consistent with differences in genome-wide mutation distribution. Some outliers also carried recurrent genomic alterations, but no single clinical, mutational or genomic feature was common to all samples or explained the persistent mutation enrichment at NFIA binding sites. This supports the NFIA-associated mutational signal as a distinct pattern that cannot be readily explained by co-binding or by a single tumour-specific process.

### Pan-cancer profiling at NFIA reveals tissue-specific mutational patterns

Since the mutational signal at NFIA binding sites was not restricted to the outlier samples (**Supplementary Fig. S5B**) and showed a mutation profile similar to that observed at CTCF binding sites in HCC samples (**Fig. 1B and 3A**), we next asked whether this pattern extended beyond liver cancer. To check this, we mapped somatic substitutions from diverse PCAWG cancer cohorts onto HepG2-defined NFIA and CTCF binding intervals (**Fig. 3G, Supplementary Fig. S7**). CTCF binding sites exhibited consistent core mutational enrichment across multiple cancer types, including melanoma and cancers of the oesophagus, bladder, breast and stomach (**Supplementary Fig. S7**), consistent with the broad susceptibility of CTCF/cohesin sites reported previously^24,45,51^. Mapping SBS to NFIA binding intervals revealed a similar, but lineage-restricted mutational landscape. While NFIA binding sites displayed significant mutation enrichment in liver (FC=1.8), kidney (FC = 1.58), prostate (FC = 1.48), pancreatic (FC = 1.23), and melanoma cohorts, they showed mutational depletion in colorectal adenocarcinoma (FC = 0.73), mirroring the known depletion of CTCF mutations in colorectal cancer particularly in DNA polymerase epsilon (POLE) mutant samples^24,60^ (FC = 0.71; **Fig. 3G, Supplementary Fig. S7**). Crucially, NFIA and CTCF mutational footprints diverged in renal and prostate carcinomas. In these malignancies, CTCF sites exhibited baseline mutation rates (FC ∼1.0), whereas NFIA sites maintained robust, significant mutational enrichment. This divergence indicates that unlike the ubiquitous, architectural mutational vulnerability of CTCF, NFIA mutational enrichment is more strongly dependent on tumour context. Along with the Mnase-seq data (**Fig. 3E**), this tissue-specific NFIA occupancy or absence of it in non-hepatic lines suggests that NFIA mutation enrichment may reflect cell-type-specific TF engagement.

### 3D chromatin architecture is associated with localised mutation accumulation

The preceding analyses examined how local TF co-binding and regulatory context were associated with mutation enrichment along the linear genome. We then considered the contribution of higher-order chromatin organisation to these patterns. To determine if 3D looping is associated with mutational accumulation across all the TFBS being assayed here, we analysed HepG2 intact Hi-C loop data from ENCODE^29^. Evaluating somatic mutation burden across loop anchors revealed a weak positive association between mutation burden and local chromatin interaction strength (Spearman ρ = 0.12; P < 1e^-4^; **Supplementary Fig. 8A and 8B**>; See Methods), suggesting that stronger loop anchors tended to harbour greater mutation accumulation.

To find out if certain TFBS were preferentially mutated within loop anchors, TFBS core regions (101 nt) were classified based on their overlap with Hi-C loop anchors, and the proportion of mutated sites was compared between loop-anchor and non-anchor TFBS (**Fig. 4A**; See Methods). Fisher’s exact test confirmed that architectural CTCF and the cohesin components RAD21 and SMC3 were significantly enriched for mutations within Hi-C loop anchors (odds ratio [OR] > 1), as expected. Multiple additional factors, including KDM6A and hepatic HNF family members, similarly displayed significant mutational enrichment at anchor-overlapping sites, although the effect was weaker than for CTCF/cohesin (**Fig. 4A, Supplementary Fig. S8D**). In contrast, mutated CEBPB binding sites were preferentially found outside loop anchors, indicating that the mutation enrichment observed for CEBPB is associated with a different genomic context (**Fig. 4A, Supplementary Fig. S8D**).

**Figure 4:**
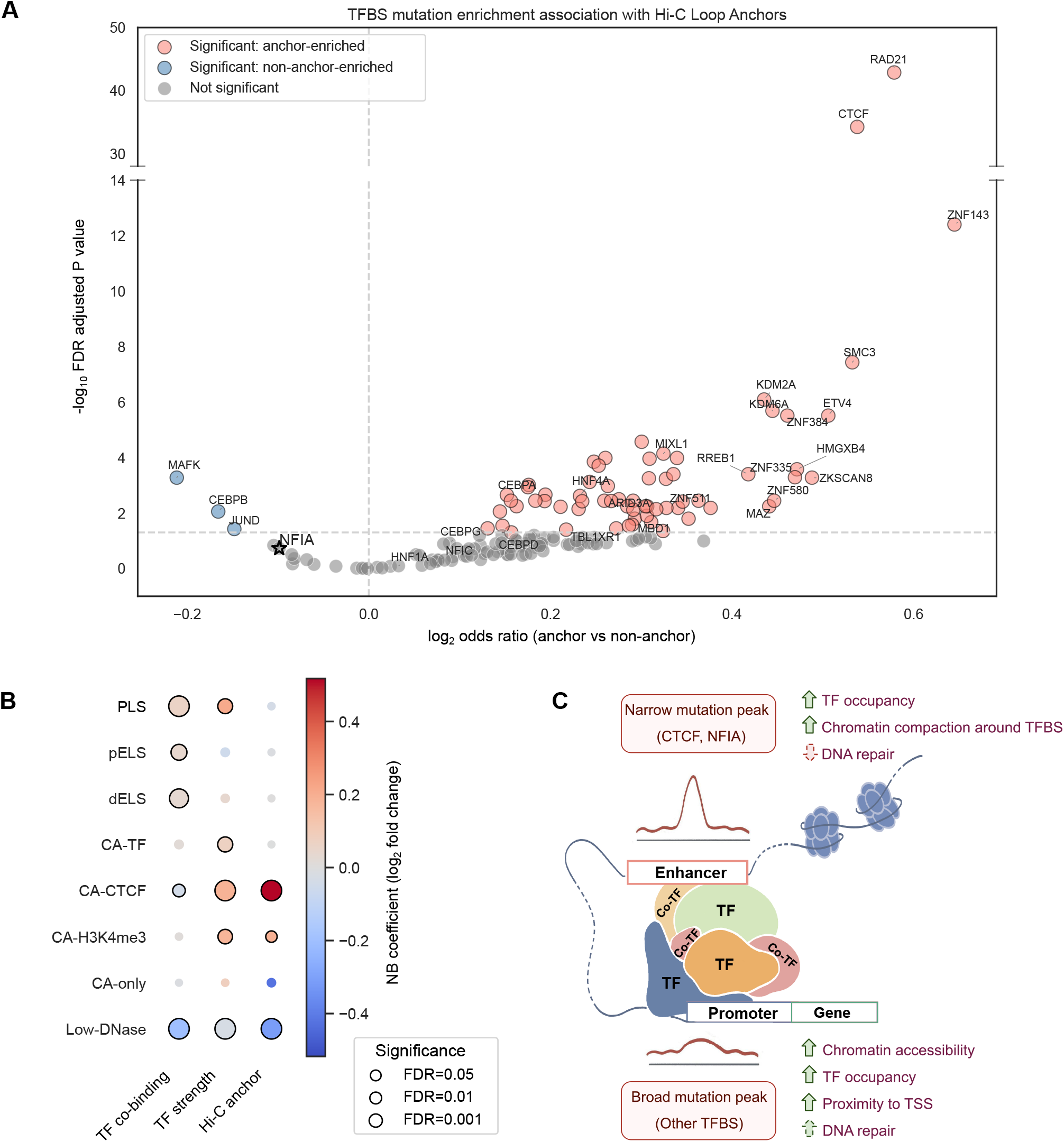
Chromatin loop architecture, cCRE features, and spatial patterns of mutational enrichment at TFBS. **A.** Differential TFBS mutational enrichment at chromatin loop anchors. Volcano plot shows the log_2_ odds ratio (OR) of mutation enrichment at TFBS within Hi-C loop anchors versus non-loop regions (on x-axis) and the corresponding significance value calculated using Fisher’s exact test followed by FDR correction (on y-axis). Positive log_2_OR indicates greater odds of mutation at TFBS overlapping loop anchors. Binding sites for the architectural factors CTCF, SMC3, and RAD21 show significant enrichment at loop anchors, whereas NFIA, highlighted by a black star, shows no loop preference. **B.** Regulatory features associated with somatic mutation burden across cCRE classes. The bubble plot shows the coefficient values (effect estimates) calculated using negative-binomial regression models fitted independently for each cCRE class. Features include the number of unique overlapping TFs, mean TF ChIP-seq signal, and Hi-C loop-anchor overlap. The colour scale represents the log2-transformed fold change from the regression model, while point size indicates FDR-adjusted significance. **C.** Schematic model of spatial mutational footprints across regulatory contexts. Summary model illustrating distinct mutational footprints across regulatory regions. Distal enhancer TFBS show high, focal mutation peaks, whereas promoter-proximal TFBS near gene bodies show broader, more diffused mutation profiles that mostly depends on cobinding contexts of TFs.

Despite harbouring high mutation rates, mutated NFIA binding sites revealed no significant preference for Hi-C loop anchors (**Fig. 4A**). Thus, unlike CTCF/cohesin-associated mutation enrichment, the elevated mutation burden at NFIA binding sites was not preferentially associated with chromatin loop anchors, suggesting that its focal enrichment is not preferentially associated with 3D architectural hubs.

### Regulatory classes show distinct associations with somatic mutation burden

Given the modest association between chromatin looping and mutation burden (prominent for CTCF/cohesin), we subsequently investigated whether other genomic regulatory features also contribute to the variation in mutation accumulation observed across TF-bound regions. Using the HepG2 candidate cis-regulatory elements (cCREs) from ENCODE SCREEN (v4)^61^, we first examined their mutation burden relative to the trinucleotide-based expectation and found most classes showed lower-than-expected mutation burden, with Chromatin Accessible-CTCF (CA-CTCF) elements showing the highest relative burden among the classes examined (**Supplementary Fig. S8C**). We further modelled mutation counts within each cCRE class using negative binomial regression (See Methods), with three features included as covariates: the number of unique overlapping TFs (n_TFs_), mean TF ChIP-seq signal (Signal_TF_), and overlap with a Hi-C loop anchor (Loop_Hi-C_). Models were fitted separately for each regulatory class, with the local trinucleotide-based expected mutation count included as an offset (See Methods). This revealed that the features associated with mutation burden differed substantially across regulatory classes (**Fig. 4B, Supplementary Fig. S8E-L**).

Consistent with our earlier findings, the strong association with 3D architecture was observed at Chromatin Accessible-CTCF (CA-CTCF) elements. Hi-C loop overlap was associated with a significantly higher mutation burden (FC = 1.43, 95% CI: 1.34 - 1.53, **Supplementary Fig. S8L**), as was the mean TF signal (FC = 1.14, 95% CI: 1.10 - 1.18). In contrast, the number of unique TFs showed a small negative association (FC = 0.97, 95% CI: 0.96 - 0.99, FDR = 0.003). Thus, mutation burden at CA-CTCF elements was most strongly associated with loop overlap and TF binding strength (**Supplementary Fig. S8L**). At regions with promoter-like signature (PLS), both TF co-occupancy and binding strength were positively associated with mutation burden (**Supplementary Fig. S8G**). Whereas at sites with enhancer-like signatures (ELS), the number of unique TFs was positively associated with mutation burden at both distal ELS and proximal ELS (**Supplementary Fig. S8H and S8J**). At CA-TF elements (Chromatin Accessible regions with TFs bound other than CTCF), mean TF signal was the only significant feature, whereas none of the three features showed a significant association at CA-only elements (that is, Chromatin Accessible regions without TF binding). At Low-DNase sites, all three features showed negative associations with mutation burden.

Together, these results show that no single feature consistently explained mutation burden across the regulatory genome. Loop overlap was most prominent at CA-CTCF elements, TF co-occupancy showed modest associations at enhancer-like elements, and TF binding strength was associated with several promoter and accessible chromatin classes. These context-dependent associations argue against a single universal driver and instead suggest that local chromatin architecture helps shape the landscape of somatic mutations across regulatory regions in cancer genomes.

## Discussion

TF binding to DNA is central to gene regulation and transcriptional control, thereby contributing to the maintenance of cellular identity and homeostasis^50^. However, persistent TF occupancy can have opposing effects on the underlying DNA, either shielding binding sites from DNA damage or, conversely, increasing their susceptibility to damage^19,26^. Previous studies of UV-induced DNA damage and repair kinetics have shown that TFBS can exhibit elevated local DNA damage, impaired repair, or both, depending on the TF family and its DNA-binding preferences^17,19,26^. CTCF and ETS-family TFs are among the most extensively characterised examples, with site-specific differences in UV-induced damage formation and repair efficiency at their binding sites contributing to elevated local somatic mutation rates in melanoma^45,51^. However, whether these effects extend beyond well-characterised TFs and tumour types, and how TF co-binding contributes to local variation in mutation rates across cancers, remain poorly understood.

In this study, we address this gap by focusing on liver cancer and leveraging one of the most comprehensive collections of ChIP-seq profiles for more than 150 TFs generated in HepG2 cells^28^. We find that somatic mutations are enriched at multiple TFBS, with motifs and core binding sites showing stronger mutational signals than their immediate flanking sequences, consistent with previous observations in melanoma^17,26^. Interestingly, our peak-shape analysis revealed that, for many TFs showing strong mutational enrichment, the signal was substantially attenuated after excluding sites co-bound by other TFs. This effect was particularly pronounced at TFBS located close to TSSs compared with those in distal regulatory regions. NFIA and CEBPB were notable exceptions, retaining significant mutational enrichment at their binding sites independently of co-binding with other TFs.

Analysis of mutation rates at TFBS in the context of chromatin-loop anchors further allowed us to distinguish effects associated with higher-order chromatin architecture from those arising from local, sequence-specific TF binding. CTCF, RAD21 and SMC3 showed greater mutational enrichment at loop anchors; however, the enrichment observed at CTCF sites was dependent on co-binding by RAD21 and SMC3, and vice versa. This finding is consistent with previous reports of elevated DNA damage and somatic mutation rates at CTCF/cohesin-binding sites^24,25,51^. In contrast, CEBPB showed significant mutational enrichment predominantly at non-looping regions, suggesting a more local and sequence-sensitive mechanism. One possible explanation is provided by observations in adult stem cells showing that C/EBP proteins can bind G:T mismatch-containing DNA with increased affinity, potentially influencing DNA repair and thereby increasing the likelihood of mutation at these sites^47^.

Similarly, mutated NFIA binding sites showed no preference for loop anchors and retained a focal mutational signal after excluding sites co-bound by other TFs. Although most NFIA-mutated sites were located distal to TSSs, the presence of active chromatin marks and bidirectional nascent transcription at these sites suggests that they reside within an active regulatory context and are likely to function as enhancers. The persistent mutation enrichment at NFIA binding sites is notable given the emerging role of NFIA in hepatocyte maturation and the broader involvement of NFI-family factors in maintaining differentiated hepatic programs^62,63^. However, whether NFIA binding itself directly influences DNA damage formation or repair remains to be determined. The depletion of specific substitution classes at the NFIA motif centre despite enrichment towards the motif edges suggests that NFIA binding may affect the directly bound motif differently from nearby DNA, for example by altering local damage formation or repair, as has been reported for other TFs^45^. Direct measurements of damage and repair at NFIA sites would be needed to test this. At the sample level, the high mutation burden observed at NFIA-binding regions was particularly evident in a subset of samples that also showed a shift towards mutations in euchromatic, early-replicating regions and exhibited recurrent CTNNB1 gains. Although CTNNB1 is a recurrently altered driver of Wnt/β-catenin signalling in HCC^59,64^, our data do not establish a direct relationship between CTNNB1 gain and NFIA-associated hypermutation. We speculate that this could be explained by the defects in DNA repair activity that may contribute to both global and local changes in mutation rate variations. The pan-cancer analysis additionally suggests that mutations at NFIA binding sites are not restricted to liver cancers but vary across tumour types. This variation is consistent with the influence of cell-of-origin chromatin on global mutation distributions^15^.

Taken together, our findings indicate that TFBS-centred mutagenesis is governed by distinct rules across different regulatory contexts. TF co-binding is associated with mutational enrichment at enhancer-like elements, whereas CTCF/cohesin-binding sites show an additional dependence on chromatin-loop architecture. NFIA represents a notable exception, retaining strong mutational enrichment at predominantly distal sites despite limited co-binding and no apparent preference for loop anchors. In contrast, promoter-proximal TFBS generally display broader and weaker mutational enrichment profiles (**Fig. 4B and 4C**). This attenuation may reflect the greater accessibility and transcription-associated DNA repair activity characteristic of active regulatory regions, although the underlying mechanisms remain unresolved^18,49^. Thus, rather than considering TFBS as a homogeneous class of genomic elements with a shared propensity for mutation, our findings suggest that TF identity, neighbouring TF occupancy, and regulatory context collectively shape their local mutational landscape. Further studies in matched cellular systems will be required to dissect the interplay between TF co-binding, DNA damage formation, and DNA repair, and to determine how these processes contribute to cancer-specific mutational patterns.

## Methods

### Datasets and quality filtering

Chromatin immunoprecipitation followed by high-throughput sequencing (ChIP-seq) data for 208 TFs and co-factors for the HepG2 cell line were retrieved from the ENCODE database^28,29^. Somatic mutation data (single-base substitutions [SBS]) for the HCC cohort (n = 314 patient tumour samples) were obtained from the PCAWG^10^ data portal. Two additional liver cohorts, 112 samples from ICGC-LICA-CN^42^ and 163 samples derived from non-malignant cirrhotic liver tissue^6^ were independently analysed.

To get enough peaks for robust mutation rate and significance analysis, we restricted the analysis to TFs with sufficient peak coverage (n ≥ 4,335 peaks, 25th percentile of the peak-count distribution), resulting in a final set of 156 (out of 208) TFs for downstream analysis. Baseline hepatic expression profiles for these 156 TFs were cross-verified using gene-level RSEM-normalized RNA-seq expression data for TCGA-LIHC and GTEx liver samples obtained from the UCSC Xena Toil RNA-seq Recompute (TcgaTargetGtex_RSEM_norm_count). Expression values are provided as log_2_(normalised count + 1)^30^. Samples were grouped as TCGA primary tumour, TCGA solid-tissue normal (adjacent/non-tumour tissue collected from cancer patients), or GTEx normal liver (donors without the corresponding cancer diagnosis) based on the accompanying phenotype annotations.

### Observed and expected mutation rate estimation

Observed somatic mutation counts were calculated by intersecting PCAWG somatic mutations (SBS) with a 2,001-bp (±1 kb) genomic window centred on the ChIP-seq peak midpoints using the pybedtools intersect (bedtools v2.31.1)^65,66^. For each TFBS, position-wise mutation counts were aggregated across their respective binding sites within the window and normalised by the total number of binding sites to derive the position-specific observed mutation rate profile.

To account for the local nucleotide composition biases, we computed the expected distribution of mutations using the genome-wide trinucleotide substitution spectrum, following approaches used previously for TFBS-centred mutation analyses^17,26^. SBS were represented by the 96 possible trinucleotide substitution contexts from the HCC cohort. For each context (c) its genome-wide mutation rate was calculated as:

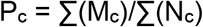

Where ∑(M_c_) represents the total number of somatic mutations occurring within the trinucleotide context (c), and ∑(N_c_) represents the total genomic occurrence of that context across the whole genome (**Fig. 1A, Supplementary Fig. S1B**). For each nucleotide position within a 2,001-bp TFBS window, the trinucleotide rate corresponding to its local sequence context was assigned as a relative mutational weight. The positional weights were normalised across the window so that they summed to one.

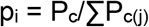

The expected mutation count at position i was then obtained by distributing the total number of observed mutations within that window, n_i_, according to these trinucleotide-derived probabilities:

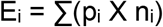

Where n_i_ is the total number of observed mutations in the window. This distributes the observed mutation count across each 2,001-bp window according to the local trinucleotide probabilities at every position, giving us the expected mutation count. These counts were subsequently aggregated across all peak regions for a given TFBS and normalised by the number of sites to yield an expected background mutation rate profile. For individual peak plots for each TF, they were further normalised by the number of samples, and are reported as mutation per megabase (Mb). Adapting the TFBS framework previously used by Frigola et al.^26^, mutational enrichment was calculated as the ratio of observed to expected mutations (FC = observed/expected) across a fixed 101-bp core region (±50-bp around the ChIP-seq peak midpoint), and statistical significance was assessed using Fisher’s exact test based on observed and expected mutation counts in the core and corresponding flanking regions, with P-values adjusted for multiple hypothesis testing using the Benjamini-Hochberg FDR procedure^67^.

### Association with ChIP-seq signal intensity

To determine how ChIP-seq signal (binding strength) of TFs at their binding site scales with mutational burden, the 156 TFBS were stratified into signal intensity tertiles: low, medium and high, based on the ChIP-seq signal values. FC was calculated independently for each tertile group using the 101-bp TFBS core. ChIP-seq signal was used as a proxy for relative occupancy of TF to DNA rather than a direct measure of binding affinity.

### Motif resolution mutational abundance

TF binding motifs within the ChIP-seq intervals were computationally identified using the position weight matrices (PWMs) from the JASPAR 2024 database (10th release)^43^ and mapped via the MOODS suite^44^(moods-dna.py -m jaspar_motif.pfm -s chip.fa -p) with a default P-value threshold for MOODS analysis (P < 10^-4^). Of the 156 TFs, high-confidence motifs were successfully mapped within the ChIP-seq regions for 90 of them (Supplementary Table 2), and their mutational profiles were precisely recentered at the JASPAR-defined motif midpoint. Both observed and expected mutation enrichment were recomputed, as described above, at the cognate motif length defined by the JASPAR model, the 101-bp core (±50-bp from the motif midpoint), the DHS-associated regions from −250 to −50-bp and +50 to +250-bp, and the remaining region was taken as distal flanking sequence within the 2,001-bp window. To assess whether the observed mutation enrichment was associated with physical TF occupancy rather than motif sequence alone, we took genome-wide TFBS motifs from JASPAR for each TF and removed regions of active binding (i.e., ChIP-seq regions to get regions with motifs but no TF binding), after which mutational enrichment was calculated as described above.

### Motif-centred substitution patterns

For mutational abundance analysis across the 17 prioritised TFBS, somatic substitutions were collapsed by strand complementarity into six pyrimidine-centred SBS classes (C>A, C>G, C>T, T>A, T>C, T>G)^68^. A fixed 41-bp window (±20-bp around the mapped motif midpoint) was used to get a high-resolution view of motif-centred substitution patterns and the immediate surrounding sequence. Positional mutation excess at each position (i) was calculated as:

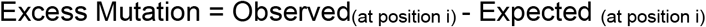

For these 17 TFBS, nucleotide composition at the 41-bp window was evaluated by aggregating all motif-containing ChIP-seq intervals. Frequencies of A, C, G and T at each coordinate were normalised (summation = 1). This was used to generate descriptive sequence-composition logos^69^.

### Peak shape (FWHM) analysis

To capture the variations in mutation peak shapes at TF bound regions, we quantified a parameter called the Full Width at Half Maximum (FWHM). For each TF, the baseline boundary of a mutational peak was defined as the genomic segment where the smoothed observed mutation profile intersected the expected mutation profile. The difference between peak maximum (Mut_max_) and base minimum (Mut_min_) within the segment was determined to obtain the height of the peak, from which half the height (H_Half_ ) of the peak was also calculated as:

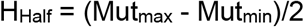

The FWHM value (in bp) is the width of the peak at half the maximum height (**Fig. 1B**). For visualisation of FWHM in bubble plots, the values across all 156 TFs were divided into four quartiles based on the complete FWHM distribution and represented using four discrete bubble sizes.

### Transcription start site proximity analysis

Gene annotation tracks were obtained from the GENCODE (v19) database^48^. To control for multiple promoters, we selected only the most upstream annotated TSS per protein-coding gene locus. Using the closest command within pybedtools^65,66^, we calculated the distance between the TFBS midpoint and the nearest TSS coordinate. Regulatory intervals falling within a window spanning 2kb upstream to 500 bp downstream of an annotated TSS (−2000 to 500-bp) were classified as proximal (promoter-associated) regions. The asymmetric window captures upstream promoter-associated regions while not extending much into the transcribed gene body. TFBS outside this defined boundary were categorised as distal regions, and the fractions of mutated TFBS classified as proximal or distal were calculated for each TF. To test whether mutation-containing TFBS differed in their TSS distribution from non-mutated TFBS, a two-sided Fisher’s exact test was performed for each TF and FDR-corrected P-values were calculated^67^.

### Pairwise TF binding overlap and Jaccard Index calculation

TF co-occupancy across the panel of 156 HepG2 factors was quantified using the Jaccard Index (JI) implemented in bedtools jaccard (v2.30.0)^66^. For any two TF binding site sets (A and B), the JI was calculated as the ratio of the intersection to the union of their ChIP-seq sites:

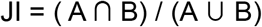

A score of 1 implies 100% similarity or overlap between the two sets, whereas 0 means that the two sets are unique and do not overlap. Pairwise JI values were compiled into a 156 X 156 similarity matrix, and co-binding modules were identified via seaborn.clustermap^70^.

### Identification of overlapping and non-overlapping TFBS

To obtain overlapping and non-overlapping binding sites from co-bound regulatory contexts, bedtools intersect through pybedtools^65,66^ was used. For a target transcription factor (x) (TF_x_), overlapping or co-bound sites were defined as TF_x_ ChIP-seq peak intervals sharing ≥ bp overlap with any peak coordinate belonging to the remaining 155 TFs, whereas non-overlapping (unique) sites were defined by removing all genomic intervals overlapping with the union of the other 155 TFs. For both overlapping and non-overlapping site subsets, observed and expected background mutation profiles, core 101-bp fold-enrichment values, and Fisher’s exact test significance were recalculated as described above.

### Leave-one-out co-binding analysis and ΔFWHM quantification

To examine whether individual co-binding partners were associated with the mutation profile of each of the 17 TFBS, a systematic leave-one-out subtraction strategy was used. For each target TF, sites overlapping one partner TF were excluded from the target TFBS set (1 vs 1). Observed and expected mutation profiles were generated as described above, and geometric peak width was evaluated at half-maximum threshold height (FWHM_after removal_). The change in peak width (ΔFWHM) was quantified as:

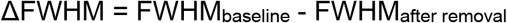

ΔFWHM matrix values across all 17 X 156 combinations were clustered using Euclidean distance metrics to resolve partner-dependency modules. Positive ΔFWHM values therefore indicate that the mutation peak became narrower after sites overlapping the partner TF were excluded, whereas negative values indicate peak broadening.

### ChIP-seq coverage, epigenomic profiling, GRO-seq, and MNase-seq analysis

To evaluate co-factor binding intensity at target TFBS, ENCODE ChIP-seq signal tracks (BigWig format) for HepG2 cells were downloaded from the ENCODE portal^28,29^. Signal coverages were extracted across a 2,001-bp window centred on the target TFBS and plotted. Histone modification datasets for hg19 included H3K9ac (GSM733638), H3K27ac (GSM733743), H3K4me3 (GSM733737), H3K9me3 (GSM1003519), H3K27me3 (GSM733754) and H3K4me1(GSM798321) fragment density BigWig tracks. HepG2 MNase-seq dyad-alignment BED files for HepG2^56^, along with processed nucleosome-density BigWig signal (MNase) tracks for K562(GSM920557) and GM12878(GSM920558), were downloaded from the ENCODE portal^29^. Histone modification profiles were generated using deepTools computeMatrix^71^, centred on NFIA binding-site midpoints across ±1 kb using 25-bp bins, and analysed separately for mutated and non-mutated NFIA binding sites. To quantify local histone-mark enrichment while excluding the TF-bound core, mean signal across the NFIA-proximal flanking regions (−500 to −100-bp and +100 to +500-bp) was compared with equal-sized local background regions (−1000 to −600-bp and +600 to +1000-bp) for each NFIA site. Proximal and background signals were compared using paired Wilcoxon signed-rank tests, with FDR corrections across histone marks and NFIA groups (Supplementary Table 3). Nascent transcriptional activity was quantified using HepG2 GRO-seq bigWig signal tracks(GSM3393700 and GSM3393701)^54^. Positive and negative-strand signals were extracted across a 2,001-bp window centred on NFIA sites and analysed separately for mutated and non-mutated TFBS.

### Outlier identification

Of the 314 samples contributing to mutation at NFIA sites, outliers were found using the interquartile range (IQR) method. Samples with total genome-wide somatic substitution counts exceeding the upper interquartile threshold (Q3 + 1.5 X IQR) were classified as cohort-wide outliers (n = 8). Samples exceeding the upper interquartile threshold for NFIA-overlapping mutations were designated as NFIA outliers (n = 13). The three samples with the highest NFIA-overlapping mutation counts were additionally analysed as the “Top 3 NFIA outliers” subset (Supplementary Table 4).

### Chromatin state mutation burden analysis

The chromatin state annotations for the HepG2 cell line were retrieved from the NIH Roadmap Epigenomics Project (ChromHMM v1.10, Core 15-state model, GRCh37)^53,57^. For this analysis, the 15 states were broadly grouped into two compartments. Euchromatin or the active state included active TSS (State 1), flanking active TSS (State 2), transcription at gene 5′/3′ regions (State 3), strong/weak transcription (States 4-5), enhancers (States 6-7), and ZNF genes/ repeats (State 8) and the inactive/heterochromatin state consisted of heterochromatin (State 9), bivalent/poised TSS (State 10), flanking bivalent TSS/enhancer (State 11), bivalent enhancer(State 12), repressed PolyComb (State 13), weak repressed Polycomb (State 14) and quiescent/low activity (State 15). Somatic mutations for each sample were intersected with these compartments using pybedtools intersect^65^ to get the observed mutations in euchromatin and heterochromatin regions. Compartment-specific observed mutation rates were calculated by normalising total observed counts to the genomic span of each compartment. Expected counts in euchromatin + heterochromatin regions were estimated using genome-wide trinucleotide substitution probabilities (P_i_). Expected counts (E_compartment_ ) were calculated by summing the product of each trinucleotide-specific mutation probability and its frequency (n_i_) within the compartment

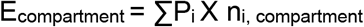

The relative compartment enrichment for each of the two compartments was quantified as Observed/Expected (O/E), and the chromatin bias per sample was expressed as a log₂ ratio of the two O/E values:

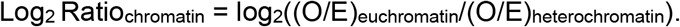

### Replication timing analysis

Genome-wide replication timing domains for HepG2 were derived from ENCODE Repli-seq datasets (GSE34399)^29,58^, using which early-replicating, transition (including both Upstream and Downstream Transition Zones [UTZ/DTZ]), and late-replicating domains were defined. Observed and expected mutation rates were computed independently for each temporal domain using the trinucleotide background model described above. Replication timing mutational bias per sample was quantified as:

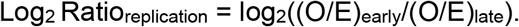

### Clinical metadata processing

Patients clinical annotations like viral infection status and donor age were retrieved from the PCAWG^10^. Normalised RNA-seq gene expression values (FPKM) for NFIA and HNF1A were extracted for annotated HCC samples^30^. Non-parametric Mann-Whitney U tests were used to evaluate the TF expression differences between virally infected (59 samples in which virus was detected) and uninfected groups (41 samples where no virus was detected). The prevalence of individual SBS signatures was compared between the 13 NFIA outlier samples and the remaining HCC samples using two-sided Fisher’s exact tests, followed by Benjamini-Hochberg FDR correction. Overall survival curves were constructed using the Kaplan-Meier analysis, and differences between NFIA outlier (n = 11/13 available) and non-outlier (n = 264/301 available) cohorts were calculated using log-rank tests. The driver gene alterations were obtained from the PCAWG patient-centric driver catalogue (Synapse syn11639581)^72^, and copy-number alterations in driver genes were identified across the NFIA outlier samples. The frequency of CTNNB1 copy-number alterations in outlier and non-outlier groups was compared using Fisher’s exact test.

### Hi-C Processing, Loop Filtering, and Anchor Metric Quantification

Processed intact Hi-C interaction datasets for the HepG2 cell line were retrieved from the ENCODE portal^29^ (GSE237899 / ENCODE accession ENCSR510MEO). Loop interaction coordinates in BEDPE format were lifted over from GRCh38 to GRCh37/hg19 using UCSC liftOver^65,73^. Only loops for which both anchors were successfully mapped were retained. As the downloaded dataset consisted of processed ENCODE loop calls, no additional filtering based on interaction enrichment, FDR or supporting pixel number was applied. For each loop, anchor lengths and midpoints were calculated from the mapped coordinates. Loop span was defined as the genomic distance between the two anchor midpoints. Somatic substitutions from the PCAWG HCC cohort were intersected with each anchor, and mutation rates were calculated from the number of overlapping substitutions relative to the corresponding anchor length and expressed as mutations per megabase. Interaction enrichment was quantified using the observed Hi-C contact signal relative to the local donut expected signal (the ring of neighbouring pixels surrounding the target contact in the Hi-C matrix) supplied with the processed loop calls:

Loop enrichment = observed contact / donut-expected contact.

Spearman rank correlation (⍴) was calculated to check association between loop enrichment and anchor mutation rate.

### TFBS mutation enrichment at Hi-C loops

To test whether mutations at individual TFBS were preferentially associated with chromatin loop anchors, 101-bp core windows centred on the ChIP-seq binding sites for each of the 156 TFs were used. Somatic substitutions from the PCAWG HCC cohort^10^ were mapped to these windows using pybedtools^65,66^. Deduplicated coordinates were generated to construct binary vectors classifying each TFBS site as either mutated or unmutated, and these were subsequently intersected with merged Hi-C anchor coordinates. For each TFBS, mutation enrichment was tested using a two-sided Fisher’s exact test based on the following four groups: mutated TFBS overlapping loop anchors (Mut_TFBS,loop_anchors_), mutated TFBS outside loop anchors ( Mut_TFBS,outside_loop_anchors_), unmutated TFBS overlapping loop anchors (NMut_TFBS,loop_anchors_), and unmutated TFBS outside loop anchors (NMut_TFBS,outside_loop_anchors_). OR was calculated as

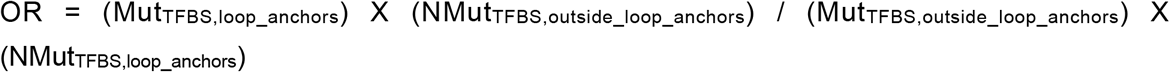

P-values were calculated using Fisher’s exact test and then adjusted for multiple testing using the Benjamini-Hochberg FDR^67^ method (FDR < 0.05).

### Full ChIP-peak analysis at Hi-C loop anchors

As a complementary analysis, the original ChIP-seq peaks of variable lengths were analysed without restricting sites to the 101-bp central window. Each peak was classified as to whether it overlapped a Hi-C loop anchor, and the somatic substitutions overlapping individual peaks were counted. For comparisons, mutation rates for loop-overlapping and non-loop-overlapping peak sets were calculated by dividing the total mutation count by the total genomic sequence represented by each peak set and expressing the resulting rate per megabase. To account explicitly for variation in individual ChIP-peak widths, mutation counts were additionally modelled using negative-binomial regression, with loop-anchor status as the predictor and the natural logarithm of peak length included as an offset. Exponentiated regression coefficients therefore represent the fold change in mutation rate for loop-anchor-overlapping peaks relative to non-overlapping peaks (**Supplementary Fig. S8D**).

### Candidate Cis-Regulatory Element (cCRE) annotation

Genome-wide candidate Cis-Regulatory Elements (cCREs, v4 release) for HepG2 were downloaded from the ENCODE SCREEN (v4) portal^61^ and lifted over to GRCh37 coordinates using the pybedtools liftOver function^65,73^. Elements were grouped into classes according to already defined ENCODE annotations: Promoter-like (PLS), Proximal Enhancers (pELS), Distal Enhancers (dELS), Chromatin Accessible CTCF (CA-CTCF), TF (CA-TF), H3K4me3 (CA-H3K4me3), Accessible only (CA-only), and Low-DNase inactive regions. We also generated a GC-matched control that did not include the cCREs plus any of the blacklisted regions from ENCODE. For each cCRE, the observed somatic mutation count was obtained by intersecting the interval with somatic substitutions from the PCAWG HCC cohort^10^. The expected mutation burden was calculated using the trinucleotide-context model described above. Briefly, context-specific mutation probabilities were summed across callable positions within each cCRE to obtain an expected mutation count. Mutation enrichment relative to this background was expressed as the ratio of observed to expected mutations (**Supplementary Fig. S8C**).

### Negative-binomial regression of cCRE mutation burden

Negative-binomial regression was used to assess the independent associations of TF occupancy, TF binding strength and Hi-C loop overlap with somatic mutation burden, consistent with previous approaches for modelling regional somatic mutation counts^74^. Models were fitted independently for each cCRE class using the statsmodels^75^ package in Python.

For each of the cCRE intervals (j) within a given class, we computed observed somatic mutation count(Y_j_). To quantify local TF occupancy, cCREs were intersected with the 156 TFs. The number of unique TFs represents the number of distinct TFs with at least one overlapping ChIP-seq peak (n_unique_TFs, j_ ). Peak signal values (binding strength) were converted to within-TF percentile ranks (from 0 to 1), and the mean signal for each cCRE was then calculated from the percentile-normalised signals of the overlapping TF peaks (Signal_j_). Both these features were z- score normalised independently within each cCRE class. Hi-C loop association was represented as a binary variable indicating whether a cCRE overlapped at least one merged HepG2 Hi-C loop anchor, using the loop-anchor dataset described above (Loop_j_). The natural log of expected mutations (E_j_) was included as an offset for the model,

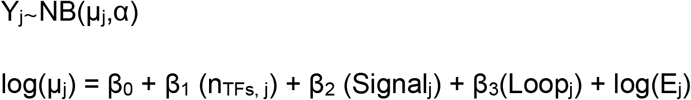

where µ_j_ represents the mean/predicted mutation count and α captures over-dispersion. Regression coefficients (β_k_), 95% confidence intervals, and P-values were extracted to assess feature impact per regulatory class. The fold change was obtained by exponentiating the regression coefficient.

## Acknowledgement

We acknowledge the funding support from the Department of Atomic Energy, Government of India, under Project Identification No. RTI 4006 and intramural funds from NCBS-TIFR. RS acknowledges support from the DBT/Wellcome Trust India Alliance Fellowship [grant number IA/ I/20/1/504928]. We thank Aswin Sai Narain Seshasayee, Deepa Agashe, and members of RS lab for their feedback and suggestions on this manuscript.

## Author contributions

NG, AKS and RS conceived and designed the study. NG gathered data, performed data analysis, interpreted the results and prepared figures. AKS provided input for the analysis and interpreted the results. NG wrote the manuscript with input from all authors. RS supervised the study. All authors read and approved the final manuscript.

## References

1. Gonzalez-Perez, A., Sabarinathan, R. & Lopez-Bigas, N. Local Determinants of the Mutational Landscape of the Human Genome. Cell 177, 101–114 (2019).

2. Alexandrov, L. B. et al. The repertoire of mutational signatures in human cancer. Nature 578, 94–101 (2020).

3. Aitken, S. J. et al. Pervasive lesion segregation shapes cancer genome evolution. Nature 583, 265–270 (2020).

4. Smith, T. C. A., Arndt, P. F. & Eyre-Walker, A. Large scale variation in the rate of germ-line de novo mutation, base composition, divergence and diversity in humans. PLOS Genet. 14, e1007254 (2018).

5. Li, R. et al. A body map of somatic mutagenesis in morphologically normal human tissues. Nature 597, 398–403 (2021).

6. Brunner, S. F. et al. Somatic mutations and clonal dynamics in healthy and cirrhotic human liver. Nature 574, 538–542 (2019).

7. Boström, M. & Larsson, E. Somatic mutation distribution across tumour cohorts provides a signal for positive selection in cancer. Nat. Commun. 13, 7023 (2022).

8. Yates, L. R. & Campbell, P. J. Evolution of the cancer genome. Nat. Rev. Genet. 13, 795–806 (2012).

9. Lawrence, M. S. et al. Mutational heterogeneity in cancer and the search for new cancer-associated genes. Nature 499, 214–218 (2013).

10. The ICGC/TCGA Pan-Cancer Analysis of Whole Genomes Consortium. Pan-cancer analysis of whole genomes. Nature 578, 82–93 (2020).

11. Hayward, N. K. et al. Whole-genome landscapes of major melanoma subtypes. Nature 545, 175–180 (2017).

12. Gröbner, S. N. et al. The landscape of genomic alterations across childhood cancers. Nature 555, 321–327 (2018).

13. Schuster-Böckler, B. & Lehner, B. Chromatin organization is a major influence on regional mutation rates in human cancer cells. Nature 488, 504–507 (2012).

14. Kikutake, C. & Suyama, M. Pan-cancer analysis of mutations in open chromatin regions and their possible association with cancer pathogenesis. Cancer Med. 11, 3902–3916 (2022).

15. Polak, P. et al. Cell-of-origin chromatin organization shapes the mutational landscape of cancer. Nature 518, 360–364 (2015).

16. Supek, F. & Lehner, B. Differential DNA mismatch repair underlies mutation rate variation across the human genome. Nature 521, 81–84 (2015).

17. Sabarinathan, R., Mularoni, L., Deu-Pons, J., Gonzalez-Perez, A. & López-Bigas, N. Nucleotide excision repair is impaired by binding of transcription factors to DNA. Nature 532, 264–267 (2016).

18. Polak, P. et al. Reduced local mutation density in regulatory DNA of cancer genomes is linked to DNA repair. Nat. Biotechnol. 32, 71–75 (2014).

19. Wasserman, H. I. et al. High-throughput characterization of transcription factors that modulate UV damage formation and repair at single-nucleotide resolution. Nat. Commun. 10.1038/s41467-026-75115-4 (2026) doi:10.1038/s41467-026-75115-4.

20. Perera, D. et al. Differential DNA repair underlies mutation hotspots at active promoters in cancer genomes. Nature 532, 259–263 (2016).

21. Morova, T. et al. Androgen receptor-binding sites are highly mutated in prostate cancer. Nat. Commun. 11, 832 (2020).

22. Yang, J. et al. Recurrent mutations at estrogen receptor binding sites alter chromatin topology and distal gene expression in breast cancer. Genome Biol. 19, 190 (2018).

23. Wong, A. M. et al. Unique molecular characteristics of NAFLD-associated liver cancer accentuate β-catenin/TNFRSF19-mediated immune evasion. J. Hepatol. 77, 410–423 (2022).

24. Katainen, R. et al. CTCF/cohesin-binding sites are frequently mutated in cancer. Nat. Genet. 47, 818–821 (2015).

25. Faseela, E. E., Notani, D. & Sabarinathan, R. CTCF/cohesin-binding sites are susceptible to replication-associated DNA damage and genomic instability in cancer cells. iScience 29, 114646 (2026).

26. Frigola, J., Sabarinathan, R., Gonzalez-Perez, A. & Lopez-Bigas, N. Variable interplay of UV-induced DNA damage and repair at transcription factor binding sites. Nucleic Acids Res. 49, 891–901 (2021).

27. Letouzé, E. et al. Mutational signatures reveal the dynamic interplay of risk factors and cellular processes during liver tumorigenesis. Nat. Commun. 8, 1315 (2017).

28. Partridge, E. C. et al. Occupancy maps of 208 chromatin-associated proteins in one human cell type. Nature 583, 720–728 (2020).

29. The ENCODE Project Consortium et al. Expanded encyclopaedias of DNA elements in the human and mouse genomes. Nature 583, 699–710 (2020).

30. Goldman, M. J. et al. Visualizing and interpreting cancer genomics data via the Xena platform. Nat. Biotechnol. 38, 675–678 (2020).

31. Sun, Y. et al. RAD21 is the core subunit of the cohesin complex involved in directing genome organization. Genome Biol. 24, 155 (2023).

32. Rao, S. S. P. et al. A 3D Map of the Human Genome at Kilobase Resolution Reveals Principles of Chromatin Looping. Cell 159, 1665–1680 (2014).

33. Odom, D. T. et al. Control of pancreas and liver gene expression by HNF transcription factors. Science 303, 1378–1381 (2004).

34. Wang, W. X., Li, M., Wu, X., Wang, Y. & Li, Z. P. HNF1 is critical for the liver-specific function of HBV enhancer II. Res. Virol. 149, 99–108 (1998).

35. Quasdorff, M. et al. A concerted action of HNF4α and HNF1α links hepatitis B virus replication to hepatocyte differentiation. Cell. Microbiol. 10, 1478–1490 (2008).

36. Fane, M., Harris, L., Smith, A. G. & Piper, M. Nuclear factor one transcription factors as epigenetic regulators in cancer. Int. J. Cancer 140, 2634–2641 (2017).

37. Chen, K.-S., Lim, J. W. C., Richards, L. J. & Bunt, J. The convergent roles of the nuclear factor I transcription factors in development and cancer. Cancer Lett. 410, 124–138 (2017).

38. Ang, L. T. et al. A Roadmap for Human Liver Differentiation from Pluripotent Stem Cells. Cell Rep. 22, 2190–2205 (2018).

39. Su, J. et al. TGF-β orchestrates fibrogenic and developmental EMTs via the RAS effector RREB1. Nature 577, 566–571 (2020).

40. Perissi, V., Aggarwal, A., Glass, C. K., Rose, D. W. & Rosenfeld, M. G. A Corepressor/ Coactivator Exchange Complex Required for Transcriptional Activation by Nuclear Receptors and Other Regulated Transcription Factors. Cell 116, 511–526 (2004).

41. Zhang, M., Huang, H., Li, J. & Wu, Q. ZNF143 deletion alters enhancer/promoter looping and CTCF/cohesin geometry. Cell Rep. 43, 113663 (2024).

42. Zhang, J. et al. International Cancer Genome Consortium Data Portal—a one-stop shop for cancer genomics data. Database J. Biol. Databases Curation 2011, bar026 (2011).

43. Rauluseviciute, I. et al. JASPAR 2024: 20th anniversary of the open-access database of transcription factor binding profiles. Nucleic Acids Res. 52, D174–D182 (2024).

44. Korhonen, J., Martinmäki, P., Pizzi, C., Rastas, P. & Ukkonen, E. MOODS: fast search for position weight matrix matches in DNA sequences. Bioinformatics 25, 3181–3182 (2009).

45. Sivapragasam, S. et al. CTCF binding modulates UV damage formation to promote mutation hot spots in melanoma. EMBO J. 40, e107795 (2021).

46. Jakobsen, J. S. et al. Temporal mapping of CEBPA and CEBPB binding during liver regeneration reveals dynamic occupancy and specific regulatory codes for homeostatic and cell cycle gene batteries. Genome Res. 23, 592–603 (2013).

47. Ershova, A. S. et al. Enhanced C/EBP binding to G·T mismatches facilitates fixation of CpG mutations in cancer and adult stem cells. Cell Rep. 35, 109221 (2021).

48. Harrow, J. et al. GENCODE: The reference human genome annotation for The ENCODE Project. Genome Res. 22, 1760–1774 (2012).

49. Zheng, C. L. et al. Transcription restores DNA repair to heterochromatin, determining regional mutation rates in cancer genomes. Cell Rep. 9, 1228–1234 (2014).

50. Lambert, S. A. et al. The Human Transcription Factors. Cell 172, 650–665 (2018).

51. Poulos, R. C. et al. Functional Mutations Form at CTCF-Cohesin Binding Sites in Melanoma Due to Uneven Nucleotide Excision Repair across the Motif. Cell Rep. 17, 2865–2872 (2016).

52. Heintzman, N. D. et al. Histone modifications at human enhancers reflect global cell-type-specific gene expression. Nature 459, 108–112 (2009).

53. Kundaje, A. et al. Integrative analysis of 111 reference human epigenomes. Nature 518, 317–330 (2015).

54. Xiao, R. et al. Pervasive Chromatin-RNA Binding Protein Interactions Enable RNA-Based Regulation of Transcription. Cell 178, 107–121.e18 (2019).

55. Core, L. J. et al. Analysis of nascent RNA identifies a unified architecture of initiation regions at mammalian promoters and enhancers. Nat. Genet. 46, 1311–1320 (2014).

56. Peng, Y. et al. Detection of new pioneer transcription factors as cell-type specific nucleosome binders. eLife 12, RP88936 (2024).

57. Ernst, J. & Kellis, M. ChromHMM: automating chromatin state discovery and characterization. Nat. Methods 9, 215–216 (2012).

58. Hansen, R. S. et al. Sequencing newly replicated DNA reveals widespread plasticity in human replication timing. Proc. Natl. Acad. Sci. U. S. A. 107, 139–144 (2010).

59. Krishna, A. et al. Mutational scanning reveals oncogenic CTNNB1 mutations have diverse effects on signaling. Nat. Genet. 58, 366–375 (2026).

60. Otlu, B. et al. Topography of mutational signatures in human cancer. Cell Rep. 42, 112930 (2023).

61. Moore, J. E. et al. An expanded registry of candidate cis-regulatory elements. Nature 10.1038/s41586-025-09909-9 (2026) doi:10.1038/s41586-025-09909-9.

62. Wesley, B. T. et al. Single-cell atlas of human liver development reveals pathways directing hepatic cell fates. Nat. Cell Biol. 24, 1487–1498 (2022).

63. Klein, K. et al. Transcription factors of the Nuclear Factor I (NFI) family control hepatocyte differentiation and cytochrome P450 activity in human liver. Pharmacol. Res. 221, 107998 (2025).

64. Schulze, K. et al. Exome sequencing of hepatocellular carcinomas identifies new mutational signatures and potential therapeutic targets. Nat. Genet. 47, 505–511 (2015).

65. Dale, R. K., Pedersen, B. S. & Quinlan, A. R. Pybedtools: a flexible Python library for manipulating genomic datasets and annotations. Bioinformatics 27, 3423–3424 (2011).

66. Quinlan, A. R. & Hall, I. M. BEDTools: a flexible suite of utilities for comparing genomic features. Bioinformatics 26, 841–842 (2010).

67. Benjamini, Y. & Hochberg, Y. Controlling the False Discovery Rate: A Practical and Powerful Approach to Multiple Testing. J. R. Stat. Soc. Ser. B Methodol. 57, 289–300 (1995).

68. Alexandrov, L. B. et al. Signatures of mutational processes in human cancer. Nature 500, 415–421 (2013).

69. Tareen, A. & Kinney, J. B. Logomaker: beautiful sequence logos in Python. Bioinformatics 36, 2272–2274 (2020).

70. Waskom, M. L. seaborn: statistical data visualization. J. Open Source Softw. 6, 3021 (2021).

71. Ramírez, F., Dündar, F., Diehl, S., Grüning, B. A. & Manke, T. deepTools: a flexible platform for exploring deep-sequencing data. Nucleic Acids Res. 42, W187–W191 (2014).

72. Sabarinathan, R. et al. The whole-genome panorama of cancer drivers. 190330 Preprint at 10.1101/190330 (2017).

73. Hinrichs, A. S. et al. The UCSC Genome Browser Database: update 2006. Nucleic Acids Res. 34, D590–D598 (2006).

74. Lee, C. A., Abd-Rabbo, D. & Reimand, J. Functional and genetic determinants of mutation rate variability in regulatory elements of cancer genomes. Genome Biol. 22, 133 (2021).

75. Seabold, S. & Perktold, J. Statsmodels: Econometric and Statistical Modeling with Python. in *Proceedings of the 9th Python in Science Conference* 92–96 (2010). doi:10.25080/Majora-92bf1922-011.

