## Supplementary Figures for "Transcription factor co-occupancy shapes the somatic mutational landscapes across regulatory regions in liver cancer"

**Supplementary Figure S1**

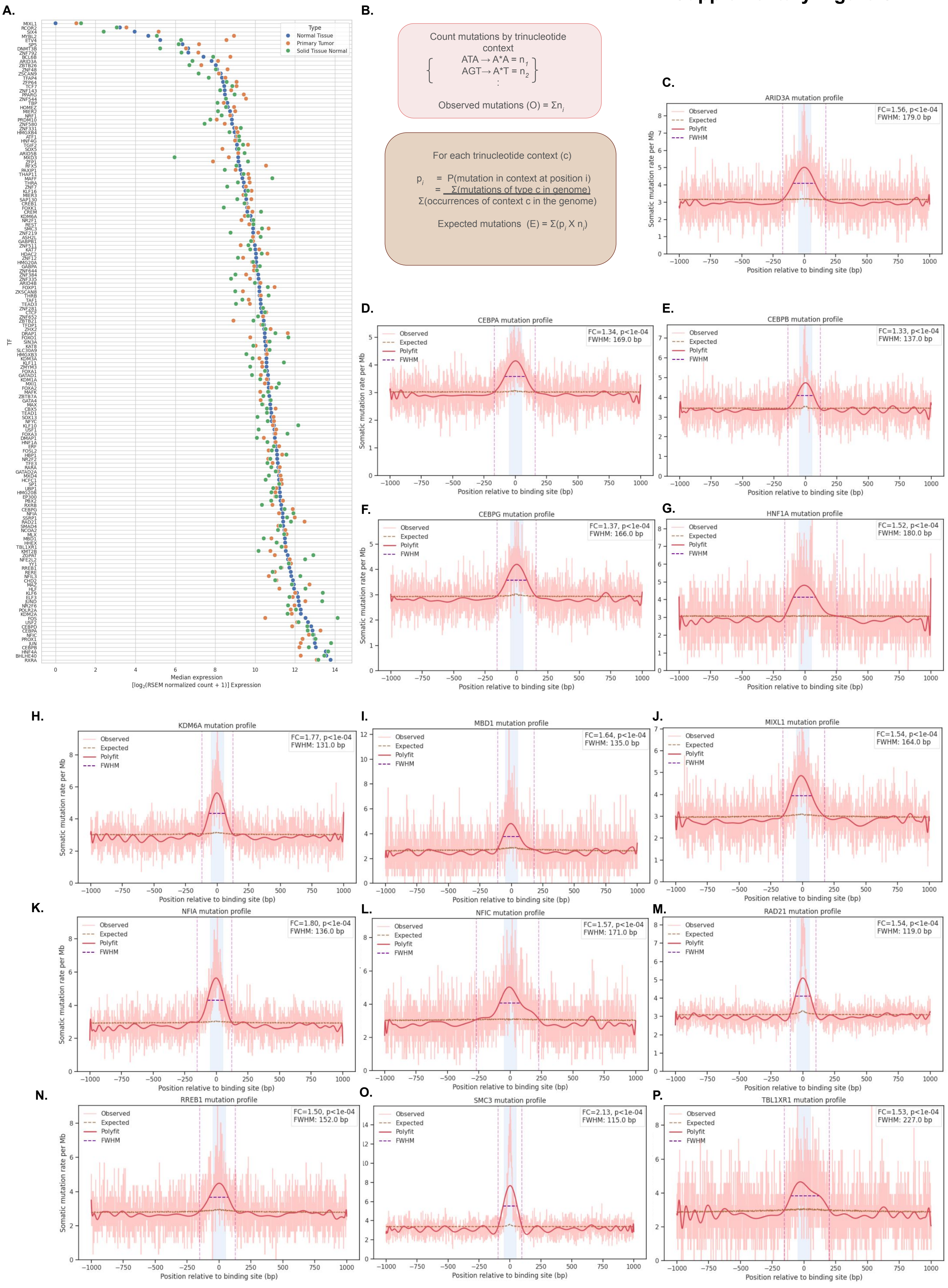

**Q.**

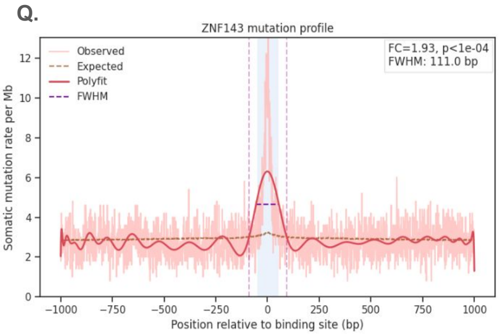


**R.**

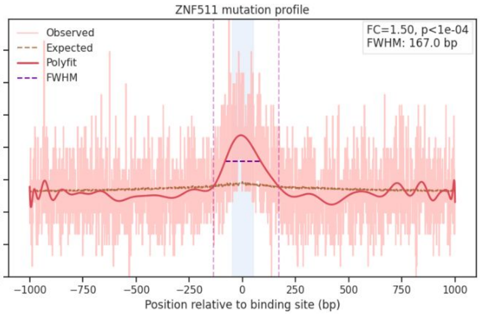


**Supplementary Figure S1: Expression of analysed TFs and mutation profiles of TFBS**

**A.** Expression levels of 156 analysed TFs across liver tissue types. Median expression levels are shown as $\log_2$(RSEM-normalised count + 1) for the 156 analysed TFs across GTEx normal liver, TCGA-LIHC solid tissue normal and TCGA-LIHC primary tumour samples.

**B.** Schematic of observed and expected mutation-rate calculation used to derive fold change (FC) across TFBS.

**C-R.** Mutation profiles for the remaining 16 prioritised TFBS: Somatic mutation rate (per Mb) across 314 samples is shown over a 2,001-bp window centred on the ChIP-seq peak midpoint. FC and statistical significance (Fisher's exact test with FDR-corrected p values) were calculated within the central 101-bp core window (±50-bp) with respect to the immediate flanking regions (excluding the core region). The red line shows the smoothed observed mutation rate, and the dashed brown line shows the expected background rate. Both core and motif significant TFBS are shown here.

**Supplementary Figure S2**

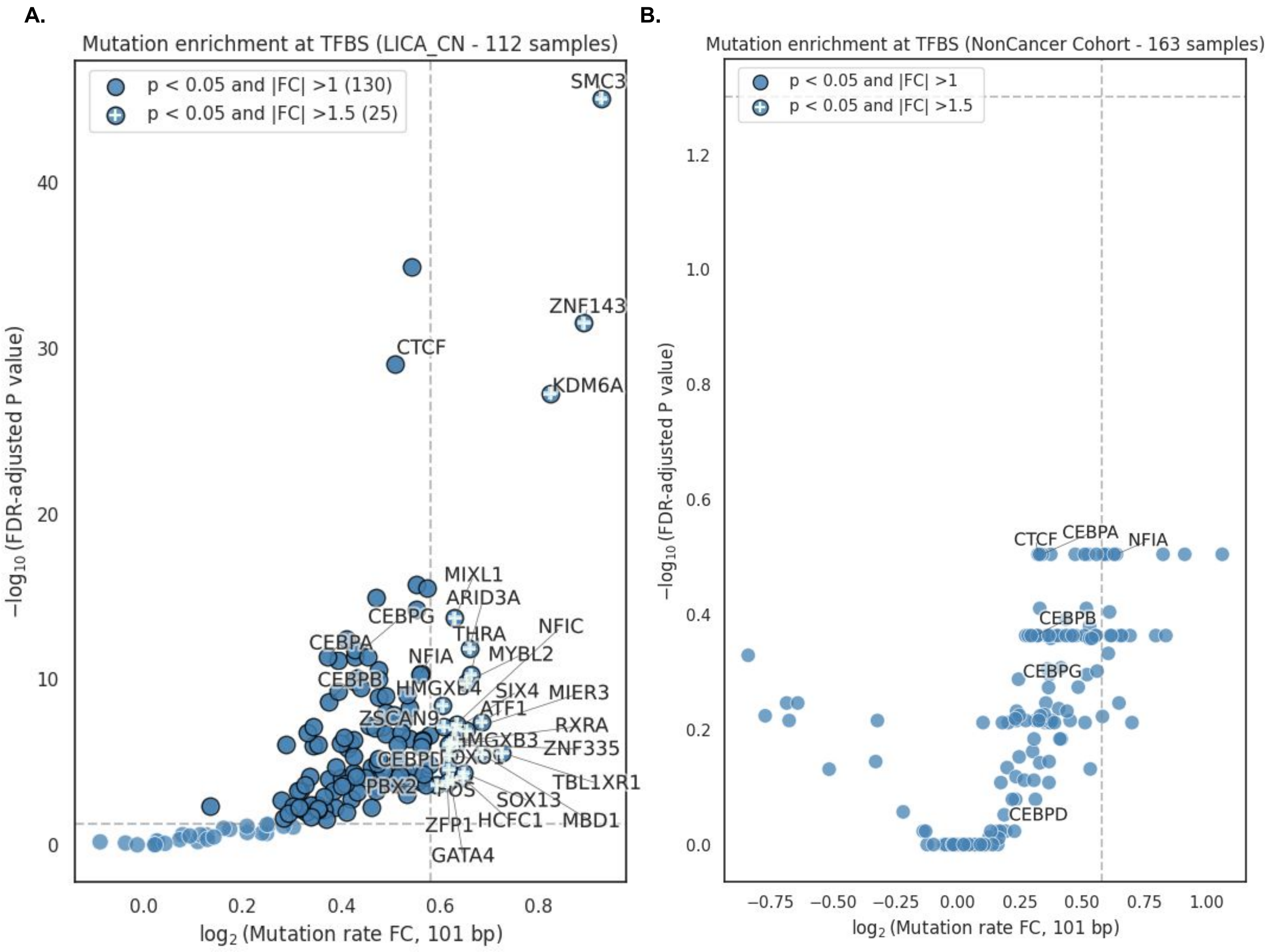


**Supplementary Figure S2: Validation of TFBS mutational enrichment in independent liver cohorts**

**A-B.** $Log_2FC$ within the 101-bp core region is plotted against $-log_{10}(FDR)$ for 156 TFs. **A.** Liver cancer (LICA-CN) **B.** Non-Cancer cohort. The horizontal dashed line marks FDR = 0.05, and the vertical dashed line marks $log_2FC$ = 0.58 (FC = 1.5). The number of significant factors is shown in brackets in the legend. Significant factors are outlined in black; significant factors with FC > 1.5 are marked "+".

### Supplementary Figure S3

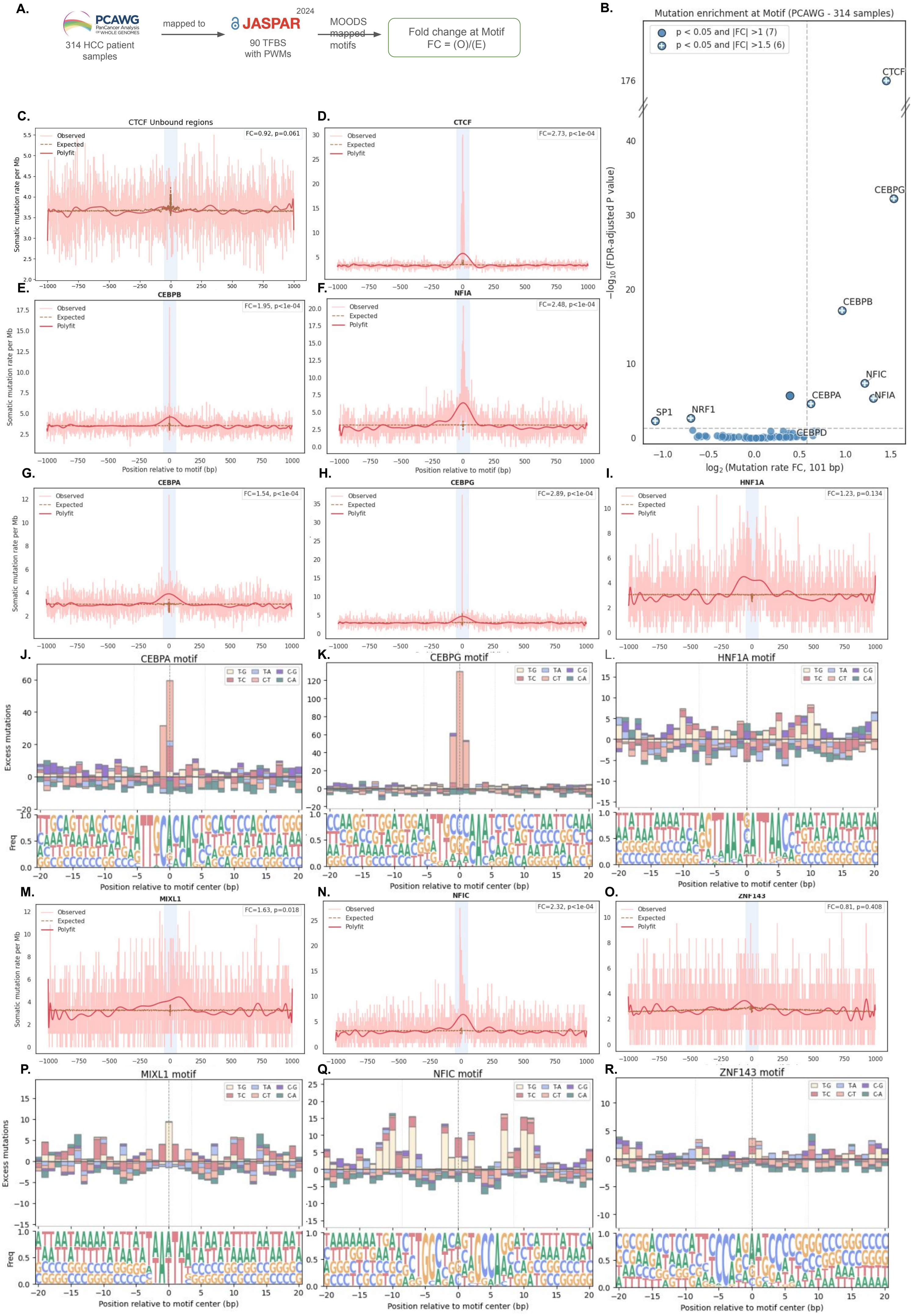

**Supplementary Figure S3: Motif-level mutation profiles at TFBS**

**A.** Schematic of motif mapping within TFBS. For factors with a corresponding JASPAR position weight matrix, motif instances were mapped within ChIP-seq peaks using MOODS to define motif-centred regions. Motif-level FC was calculated from mutations overlapping the mapped motif instances.

**B.** Mutational enrichment across 90 TFs with mapped motifs. $Log_2FC$ at the TF motif is plotted against $-log_{10}(FDR)$ for 90 TFs in the PCAWG-HCC cohort. The horizontal dashed line marks FDR = 0.05, and the vertical dashed line marks $log_2FC = 0.58$ (FC = 1.5). The number of significant factors is shown in brackets in the legend. Significant factors are outlined in black; significant factors with FC > 1.5 are marked “+”.

**C.** Mutation profile at unoccupied CTCF motif instances is shown as a representative example. Somatic mutation rate (per Mb) is shown across a 2,001-bp window centred on JASPAR CTCF motifs that do not overlap HepG2 CTCF ChIP-seq peaks, serving as an unoccupied motif control.

**D-F.** Mutation profiles at bound TFBS motifs. Somatic mutation rate (per Mb) is shown across a 2,001-bp window centred on mapped motif midpoint for **(D)** CTCF , **(E)** CEBPB and **(F)** NFIA ChIP-seq peaks. The red line shows the smoothed observed mutation rate, and the dashed brown line shows the expected background rate.

**G–I and M–O.** Motif-centred mutation-rate profiles for CEBPA **(G)**, CEBPG **(H)**, HNF1A **(I)**, MIXL1 **(M)**, NFIC **(N)** and ZNF143 **(O)**. Somatic mutation rate (per Mb) is shown across a 2,001-bp window centred on the mapped ChIP-seq motif midpoint. The red line shows the smoothed observed mutation rate and the dashed brown line the expected background rate.

**J–L and P–R.** Corresponding motif-centred substitution profiles and nucleotide frequency logos for CEBPA **(J)**, CEBPG **(K)**, HNF1A **(L)**, MIXL1 **(P),** NFIC **(Q)** and ZNF143 **(R)**. The upper panels show observed-minus-expected mutations across pyrimidine-centred substitution classes within a 41-bp motif-centred window; the lower panels show nucleotide frequency logos.

**Supplementary Figure S4**

A.

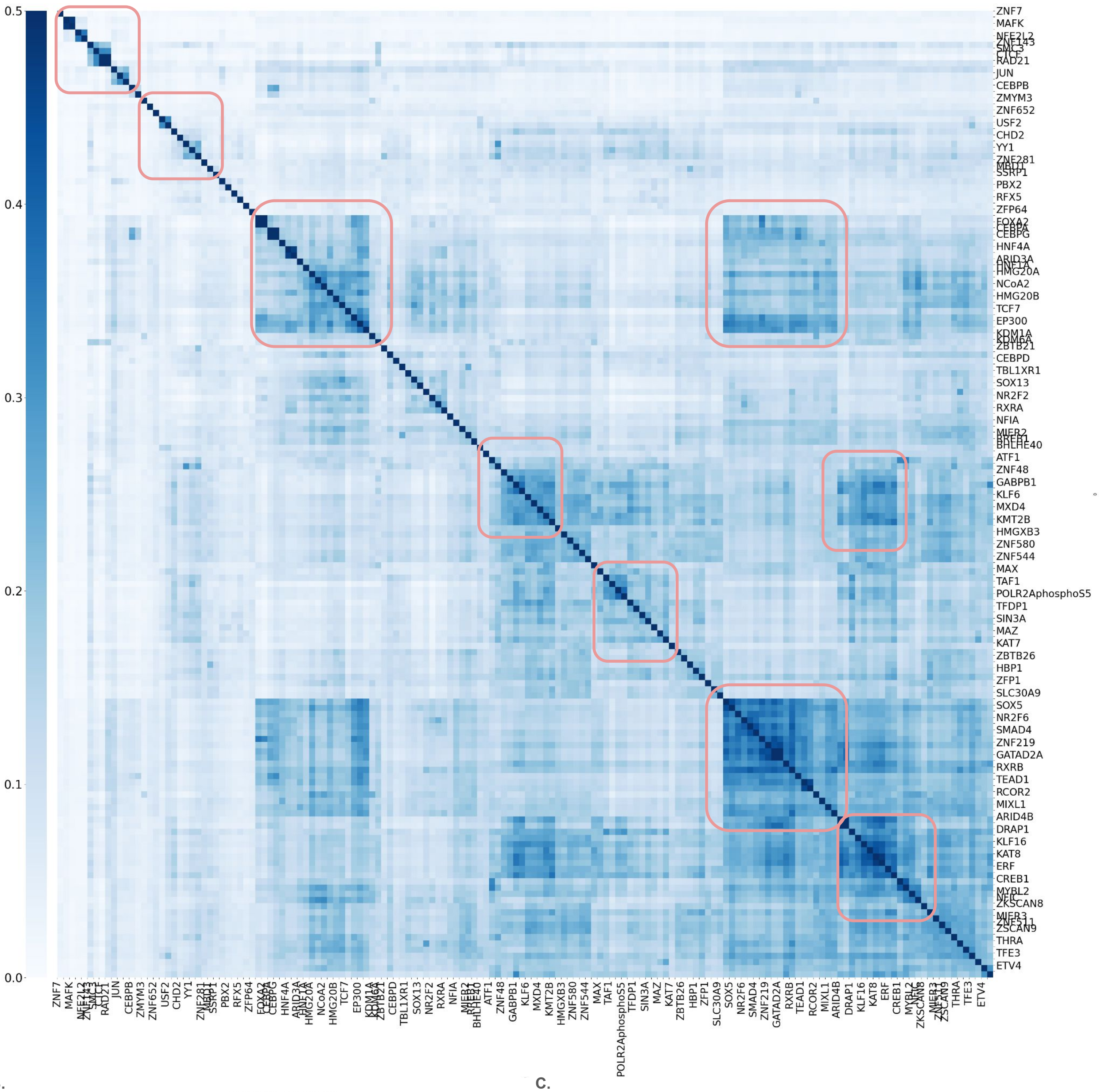


B.

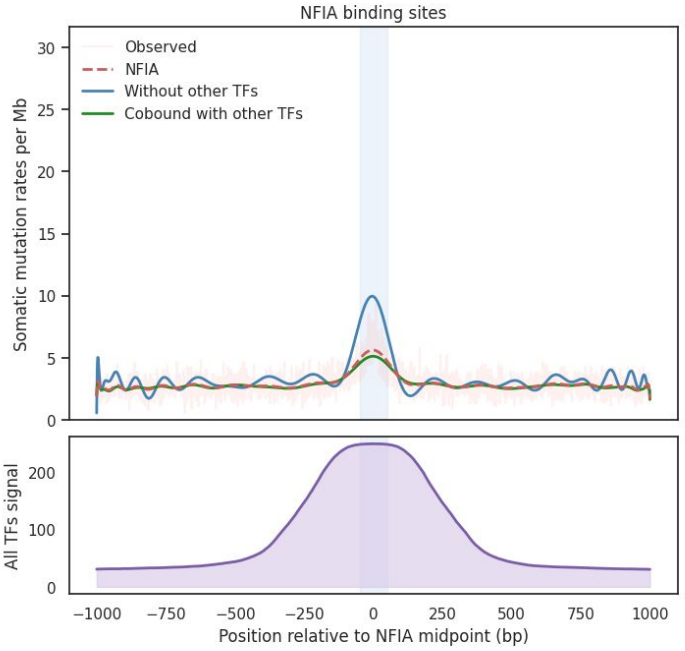


C.

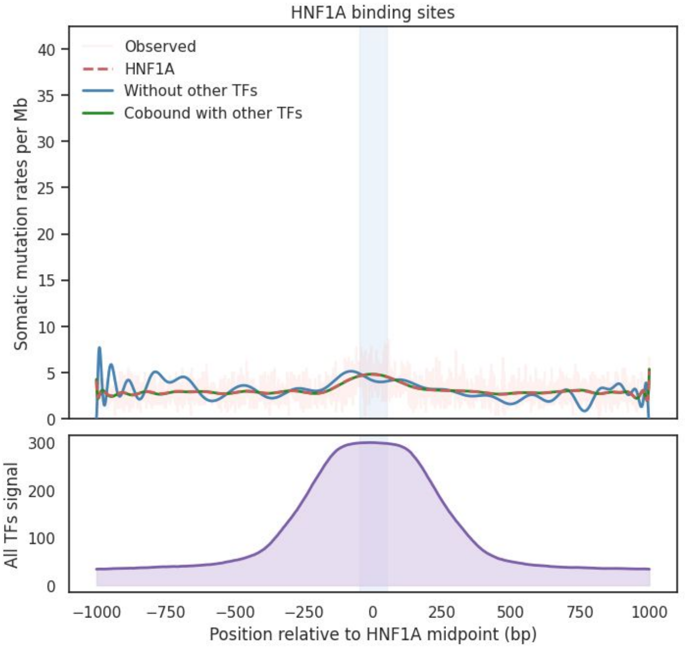

**Supplementary Figure S4: TF co-binding clusters and mutation profiles after removal of overlapping TFBS**

**A.** Co-binding similarity across TFBS. Jaccard similarity (Jaccard Index -JI) scores were calculated for all pairwise combinations of the 156 TFBS sets based on overlap between binding sites. Hierarchical clustering of the resulting 156 × 156 similarity matrix identifies multiple TFBS overlap modules (outlines in red). The bar on the left represents the JI range for the heatmap.

**B.** Mutational footprints and cofactor occupancy at NFIA binding sites. The upper panel shows somatic mutation rate profiles (per Mb) across a 2,001-bp window at all NFIA binding sites (FC = 1.80, $p < 1e\text{-}04$), NFIA sites after removing sites overlapping any of the other 155 TFBS sets (FC = 2.73, $p < 1e\text{-}04$), and NFIA sites co-bound by at least one other TF (FC = 1.67, $p < 1e\text{-}04$). The lower panel shows aggregate ChIP-seq coverage for the remaining 155 TFs across the same NFIA-centred window.

**C.** Mutational footprints and cofactor occupancy at HNF1A binding sites. The upper panel shows somatic mutation rate profiles (per Mb) across a 2,001-bp window at all HNF1A binding sites (FC = 1.52, $p < 1e\text{-}04$), HNF1A sites after removing sites overlapping any of the other 155 TFBS sets (FC = 1.48, $p = 0.163$), and HNF1A sites co-bound by at least one other TF (FC = 1.52, $p < 1e\text{-}04$). The lower panel shows aggregate ChIP-seq coverage for the remaining 155 TFs across the same HNF1A-centred window.

**Supplementary Figure S5**

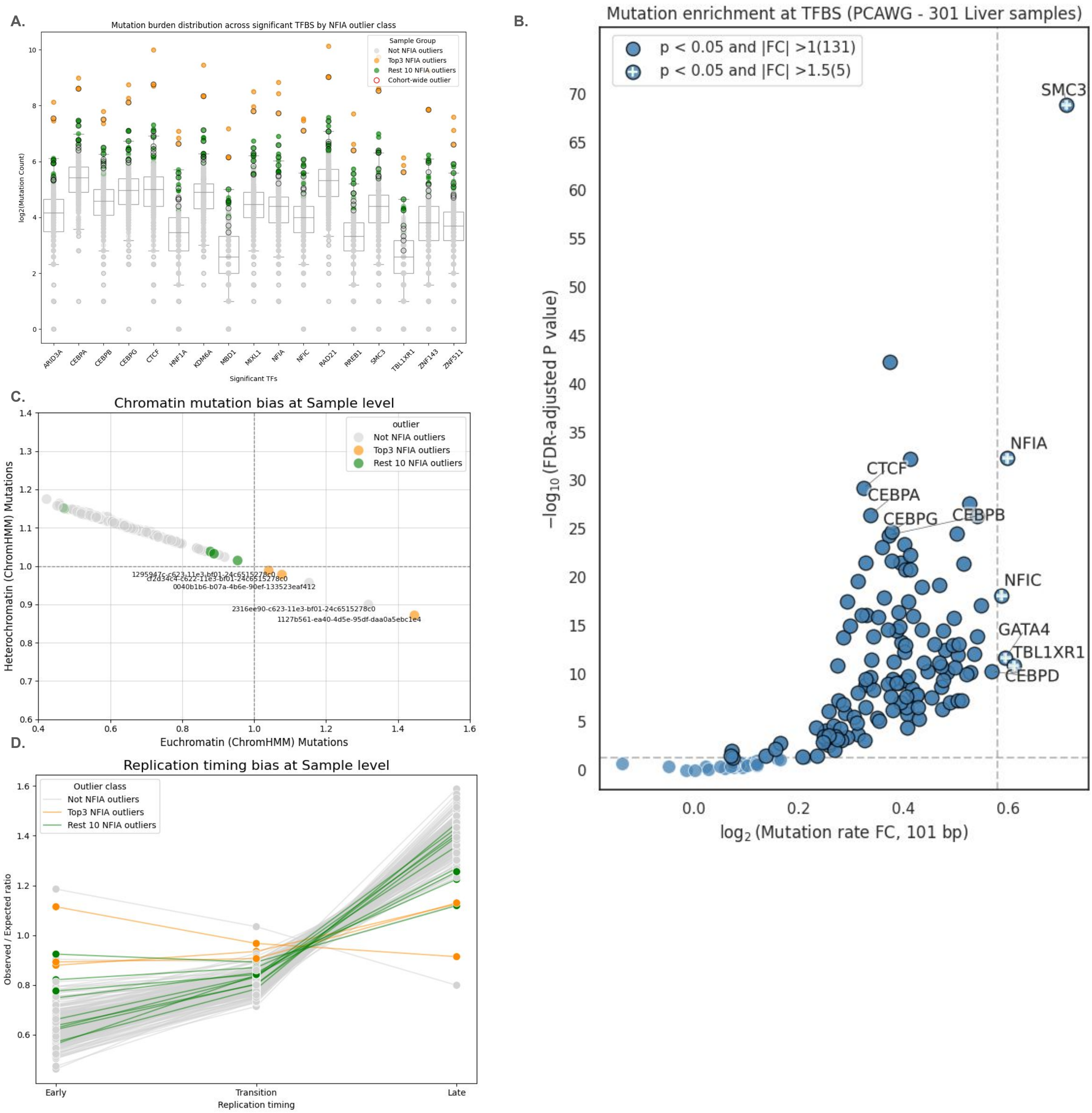

**Supplementary Figure S5: Mutation and genomic features of NFIA mutation-burden outlier samples**

**A.** Sample-wise mutation burden across significant TFBS. Box plot showing $\log_2$ mutation counts across PCAWG liver cancer samples at 17 prioritised TFBS. NFIA mutation-burden outliers were identified using the IQR method and show elevated mutation burden across multiple TFBS.

**B.** TFBS mutational enrichment after outlier exclusion. $\log_2 FC$ within the 101-bp core region is plotted against $-\log_{10}(FDR)$ for 156 TFBS after excluding the 13 NFIA mutation-burden outliers. The horizontal dashed line marks FDR = 0.05, and the vertical dashed line marks $\log_2 FC = 0.58$ (FC = 1.5). The number of significant factors is shown in brackets in the legend. Significant factors are outlined in black; significant factors with FC > 1.5 are marked "+".

**C.** Sample-level mutation burden across chromatin states. Scatter plot comparing sample-wise mutation enrichment (observed/expected) in active and repressed chromatin states defined using HepG2 ChromHMM annotations. The top three NFIA outliers, highlighted in orange, show greater relative mutation enrichment in active chromatin.

**D.** Mutation rates across replication-timing domains. Mutation enrichments (observed/expected) are shown across early, transition and late replication-timing domains in HepG2 cells, comparing the full cohort with the top NFIA outlier samples.

**Supplementary Figure S6**

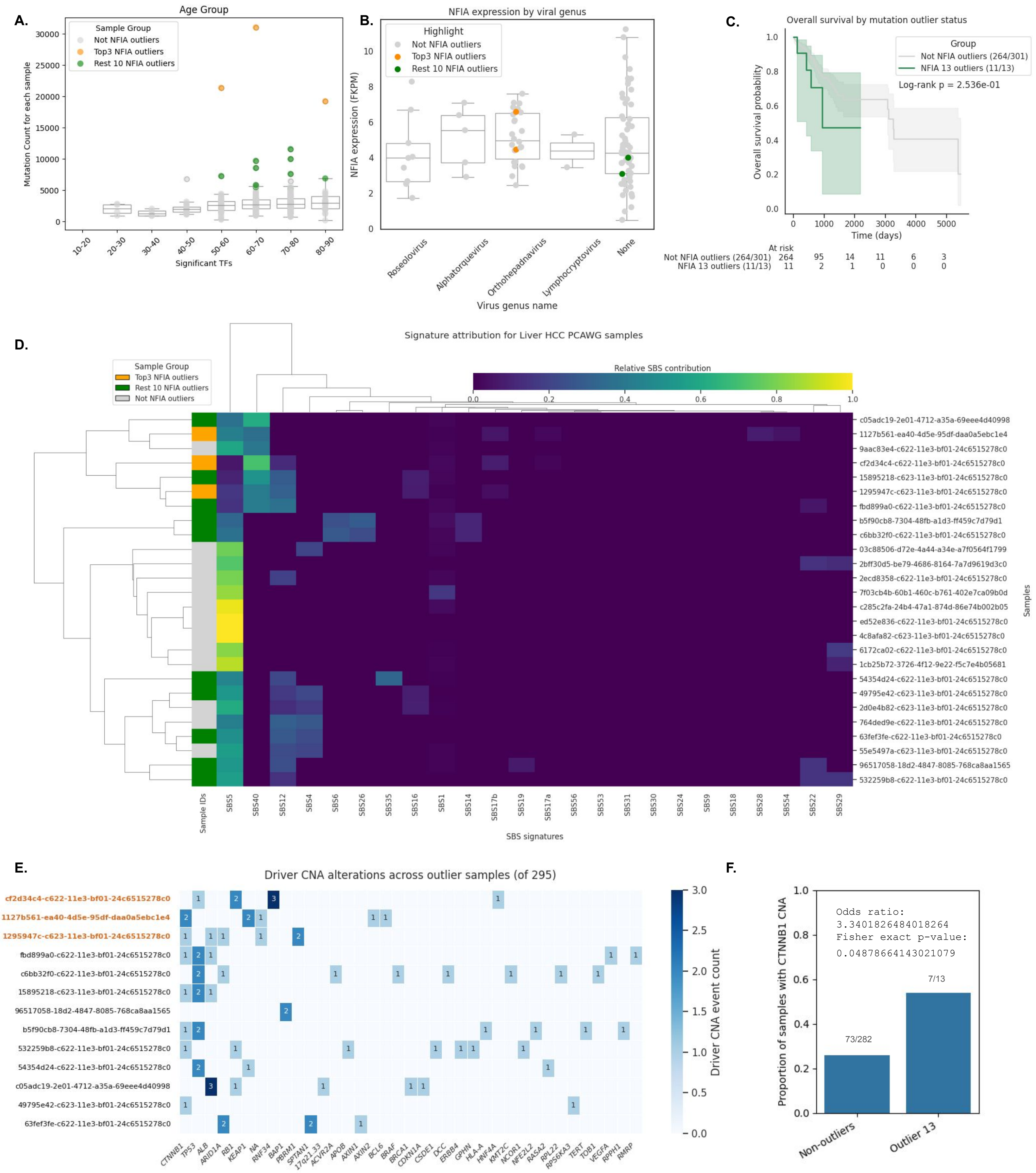

**Supplementary Figure S6: Clinical, mutational signature, and driver alterations associated with NFIA outlier samples**

**A.** Patient age across cohort subsets. Box plot showing donor age distributions between NFIA outliers and non-outlier samples.

**B.** NFIA expression by viral infection status. Box plot comparing NFIA gene expression across HCC samples grouped by hepatitis viral infection status and uninfected controls. Significance was evaluated using the Mann–Whitney U test.

**C.** Overall survival of NFIA outlier patients. Kaplan–Meier curves comparing overall survival between NFIA outliers and non-outlier samples. Significance was evaluated using the log-rank test.

**D.** Mutational signature profiles of NFIA outliers. Clustered heatmap showing SBS signature contributions across the 13 NFIA outlier and 13 randomly selected non-outlier samples.

**E.** Copy-number alterations in driver genes across NFIA outliers. Discrete heatmap showing copy-number alterations affecting PCAWG patient-centric driver genes across the NFIA outlier samples, including recurrent alterations at the CTNNB1 locus.

**F.** Proportion of CTNNB1 copy-number alterations in outlier and non-outlier groups. Significance was evaluated using Fisher's exact test.

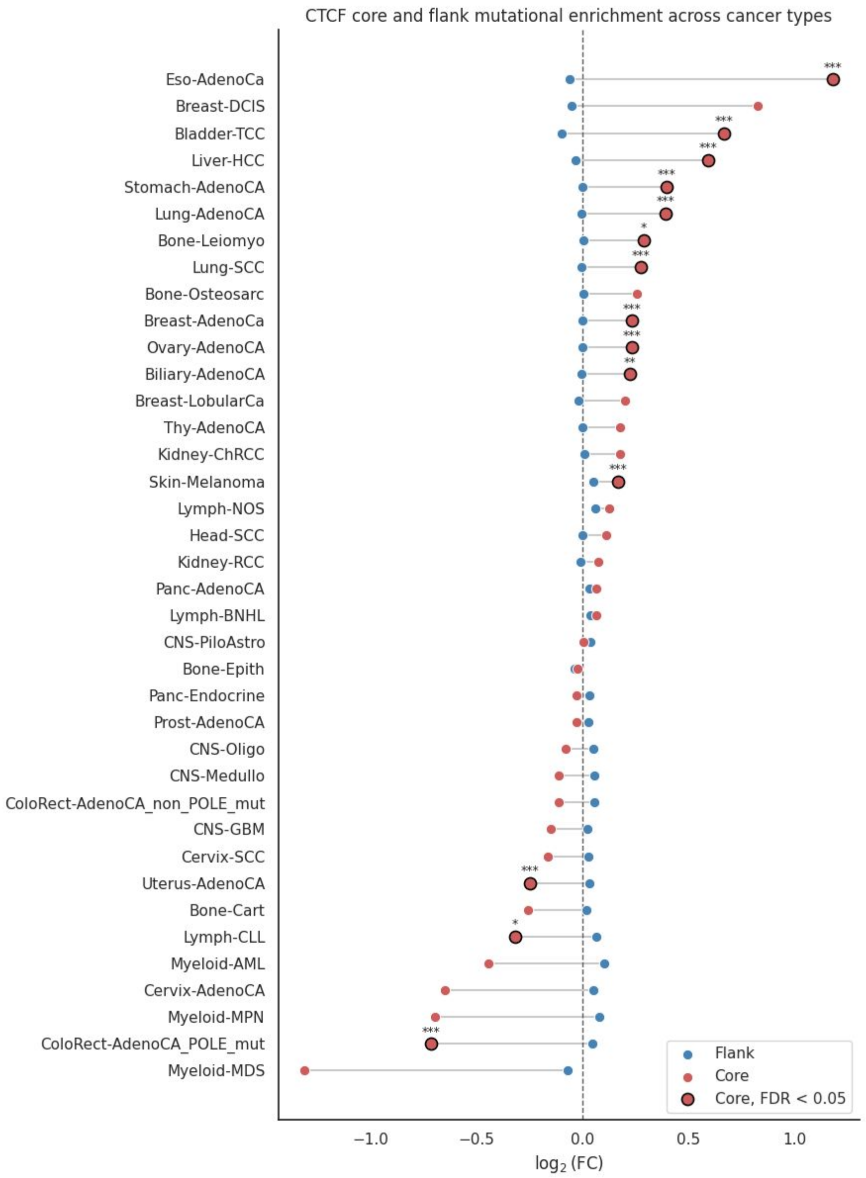


**Supplementary Figure S7: Pan-cancer mutational enrichment at HepG2-defined CTCF binding sites**

Somatic substitutions from individual PCAWG cancer cohorts were mapped to CTCF binding intervals defined in HepG2 cells. $Log_2FC$ is shown for the central 101-bp core and flanking regions for each cancer cohort. Significant enrichment or depletion determined by Fisher's exact test is denoted as *P < 0.05, **P < 0.01 and ***P < 0.001.

### Supplementary Figure S8

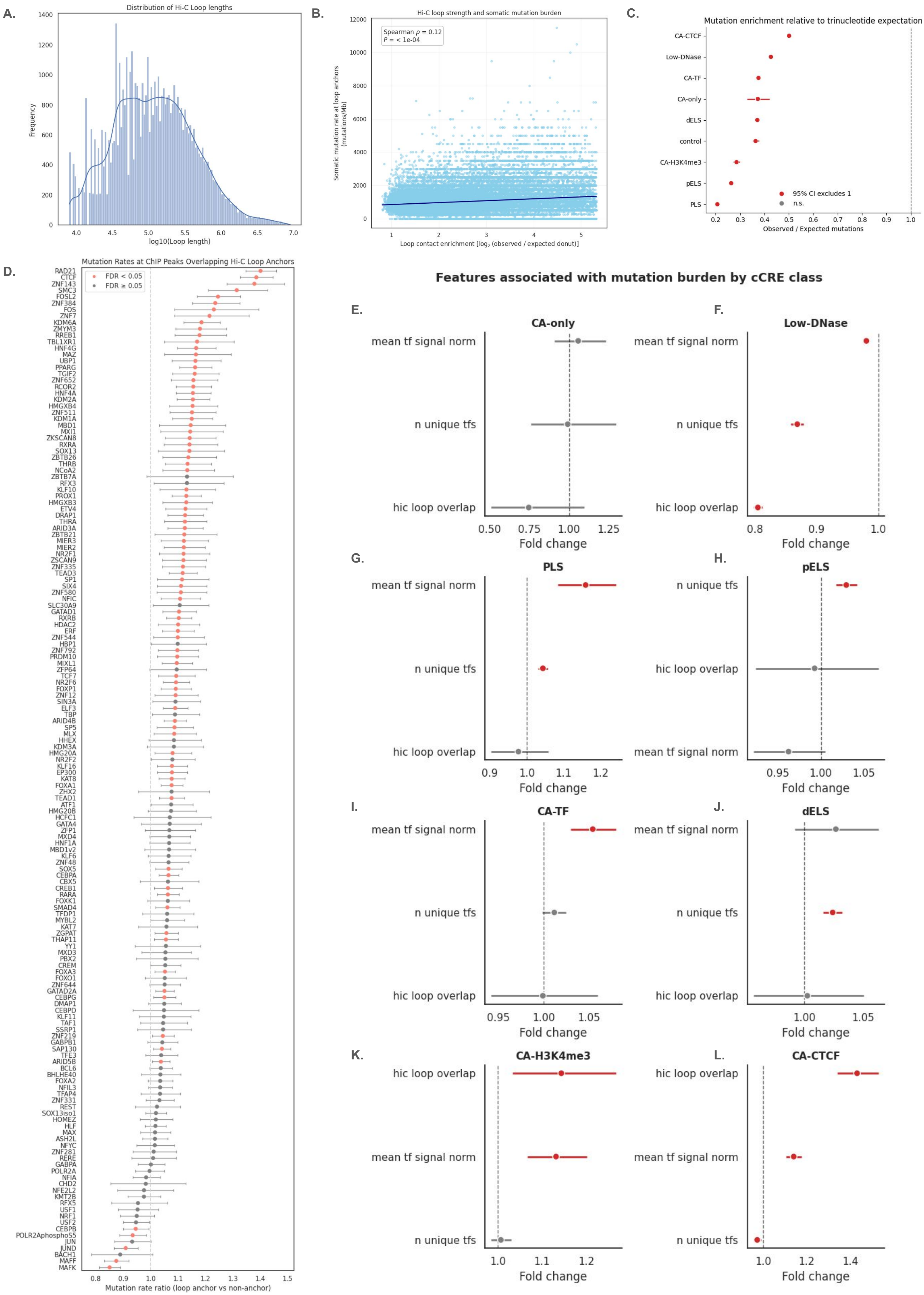

**Supplementary Figure S8: Chromatin loop properties and genomic features associated with cCRE mutation burden**

**A.** Distribution of HepG2 Hi-C loop spans. Distribution of genomic distances between the midpoints of the two interacting loop anchors, shown on a $\log_{10}$ scale.

**B.** Association between loop enrichment and mutation rate at chromatin loop anchors. Scatter plot showing somatic mutation rates per megabase at loop anchors as a function of Hi-C loop enrichment. Association was assessed using Spearman rank correlation.

**C.** Somatic mutation enrichment across cCRE classes. Mutation enrichment is shown as the ratio of observed mutations to the trinucleotide-based expected mutation burden for each cCRE class. A value of 1 indicates that the observed mutation burden equals the expected burden.

**D.** Mutation enrichment at ChIP-seq peaks overlapping Hi-C loop anchors. Forest plot showing the estimated FC in mutation rate for full ChIP-seq peaks overlapping loop anchors compared with non-overlapping peaks. Effects were estimated using negative-binomial regression while accounting for differences in peak length. Significant associations are highlighted in red, and horizontal error bars indicate 95% confidence intervals.

**E-L.** Genomic features associated with mutation burden across cCRE classes. Exponentiated negative-binomial regression coefficients (fold changes) showing the associations of the number of unique overlapping TFs, mean TF ChIP-seq signal, and Hi-C loop-anchor overlap with mutation burden within each cCRE class. Horizontal error bars indicate 95% confidence intervals.
